# A non-olfactory shark adenosine receptor activates CFTR with unique pharmacology and structural features

**DOI:** 10.1101/2020.11.01.363762

**Authors:** Sumeet Bhanot, Gabriele Hemminger, Cole L. Martin, Stephen G. Aller, John N. Forrest

## Abstract

Adenosine receptors (ADORs) are G-protein coupled purinoceptors that have several functions including regulation of chloride secretion via CFTR in human airway and kidney. We cloned an ADOR from *Squalus acanthias* (shark) that likely regulates CFTR in the rectal gland. Phylogenic- and expression- analyses indicate that elasmobranch ADORs are non-olfactory, and appear to represent extant predecessors of mammalian ADORs. We therefore designate the shark ADOR as the A_0_ receptor. We co-expressed A_0_ with CFTR in *Xenopus laevis* oocytes and characterized the coupling of A_0_ to the chloride channel. Two electrode voltage clamping was performed and current-voltage (I-V) responses were recorded to monitor CFTR status. Only in A_0_- and CFTR- co-injected oocytes did adenosine analogs produce a significant concentration-dependent activation of CFTR consistent with its electrophysiological signature. A pharmacological profile for A_0_ was obtained for ADOR agonists and antagonists that differed markedly from all mammalian ADOR subtypes (agonists: R-PIA > S-PIA > CGS21680 > CPA > 2ClADO > CV1808 = DPMA > NECA) and (antagonists: DPCPX > PD115199 > 8PT > CGC > CGS15943). Structures of human ADORs permitted a high-confidence homology model of the shark A_0_ core which revealed unique structural features of ancestral receptors. We conclude: (1) A_0_ is a novel and unique adenosine receptor ancestor by functional and structural criteria; (2) A_0_ likely activates CFTR *in vivo* and this receptor activates CFTR in oocytes indicating an evolutionary coupling between ADORs and chloride secretion; and (3) A_0_ appears to be a non-olfactory evolutionary ancestor of all four mammalian ADOR subtypes.

**Significance Statement:** We have cloned and characterized an ancient adenosine receptor from sharks that is unlikely to be olfactory in function. The shark receptor, which we designate as A_0_, has a unique pharmacological profile, characteristic structural features, and is also highly likely to be the dominant ADOR regulator of the shark ancient ortholog of the Cystic Fibrosis chloride channel, called CFTR.

## Introduction

Two broad classes of purinoceptors, P_1_ & P_2_, recognize and bind adenosine-based nucleosides and nucleotides, and can be distinguished by two anti-parallel pharmacological profiles (P_1_: Adenosine > AMP > ADP > ATP & P_2_: ATP > ADP > AMP > Adenosine) (1). Receptors specific for adenosine (ADORs) are divided into four non-olfactory subtypes including A_1_, A_2a_, A_2b_ & A_3_, which couple to G_o,i(1-3)_, G_s_, G_s_ and G_i(2,3),q-like_ G proteins, respectively (2–7). Typically, all ADORs modulate the activity of at least one of a number of different isoforms of adenylyl cyclase through the coupling to a G protein. The A_1_ ADOR can also modulate other effector systems, namely potassium channels, calcium channels, phospholipase A and C (7).

One major unifying theme in the function of ADORs in mammals appears to be a feedback mechanism to either curb metabolic activity of the cell or to oppositely vasodilate for increasing oxygen supply and meet higher metabolic demand (8). Relatedly, in sharks, which evolved prior to any land-based vertebrate, low concentrations of adenosine released from metabolic activity by the highly active shark rectal gland of *Squalus acanthias*, serve as an autocoid feedback inhibitor to slow ion transport (9). Curiously, high concentrations of adenosine (~10 μM) increased ion transport activity at both the apical- and basolateral-membranes, and stimulated chloride secretion in rectal gland via the shark ortholog of the Cystic Fibrosis transmembrane Conductance Regulator (CFTR) (10,11). Shark CFTR is an impressive 70% identical to human CFTR at the amino acid residue sequence level (12), and an ancient evolutionary link in ADOR signaling of CFTR activity in both fish and mammals is now well established.

The four main ADOR subtypes (A_1_, A_2a_, A_2b_ & A_3_), first identified in many different mammals and now observed in the genomes of birds, reptiles, amphibians and fish, can be distinguished at phylogenetic- and pharmacological levels. For the A_1_ receptor, the adenosine analogs CPA and R-PIA are the most potent, while CV1808 and CGS21680 are weaker agonists. For A_2a_, NECA, CV1808, and CGS21680 are the most potent agonists, and CPA is less potent. The A_2b_ receptor subtype responds with high potency to NECA but exhibits little or no response to CV1808 and CGS21680. A fifth ADOR subtype, designated A2c, was more recently identified and was shown to be an olfactory ADOR that allows fish to sense adenosine released into the water by potential prey (13).

We sought to examine ADORs in *Squalus acanthias* and we identified a single receptor with highest expression in the rectal gland and stomach, but very low levels in brain and other tissues. Phylogenetic analysis shows that the *Squalus acanthias* ADOR, as well as putative ADORs identified by genome sequencing from other sharks, amphibians and reptiles, form a separate clade that is distinct from the other five ADOR subtypes. We used the *Xenopus laevis* oocyte system to co-express the *Squalus acanthias* ADOR with CFTR to demonstrate the coupling of receptor to chloride channel. Co-injected oocytes revealed a pharmacological profile of *Squalus acanthias* ADOR that is unique from all four major subtypes of ADORs. We argue that the *Squalus acanthias* ADOR is not olfactory and that the clade containing the elasmobranch ADORs is ancestral to the four major subtypes. We therefore designated the shark ADOR as A_0_-ADOR, and characterize this ancestral receptor here including novel structural features revealed by three-dimensional atomic homology modeling.

## Materials and Methods

### Molecular Biology

Tissues were obtained from male dogfish sharks, *Squalus acanthias*, weighing 2-4 kg, which were caught by gill nets in Frenchman’s Bay, ME, and kept in marine live cars until use, usually within 3 days of capture. One male dogfish shark was killed by pithing the spinal cord. Details describing the molecular biology (preparation of total RNA from shark tissue, degenerate PCR, library screening, RACE PCR, Northern blot, qRT-PCR and electrophysiology in oocytes) are presented in the supplementary material (see supplementary figs. S1-S5, https://www.biorxiv.org/content/biorxiv/early/2020/11/02/2020.11.01.363762/DC1/embed/media-1.pdf?download=true). Human CFTR was obtained from Dr. Carol Semrad in the pKS-βglobin-CFTR-polyAC plasmid and human adenosine receptors were provided by Dr. Joel Linden. Capped messenger RNA for all receptors was synthesized from linearized plasmid templates using the Ambion mMessage mMachine T7 *in vitro* transcription kit according to the manufacturer’s protocol.

To retrieve putative adenosine receptor coding sequences with good representation from mammals, birds, reptiles, amphibians and fish, multiple database searches were carried out using the NCBI BlastP Server against the full nonredundant dataset. To find potential ancestral receptor sequences, additional searches using shark A_0_ sequence were performed excluding the Mammalia taxid (40674). Global alignments were performed using the Clustal method (14) within the Lasergene software package (DNASTAR, Inc) and similar results were obtained using the implementation of Clustal Omega (15) in the EMBOSS package (16) and standard parameters (17). Alignments for phylogenetic analyses including the production of the phylogenetic tree in figure 1 were conducted with MEGA7 (18).

**Figure 1.**
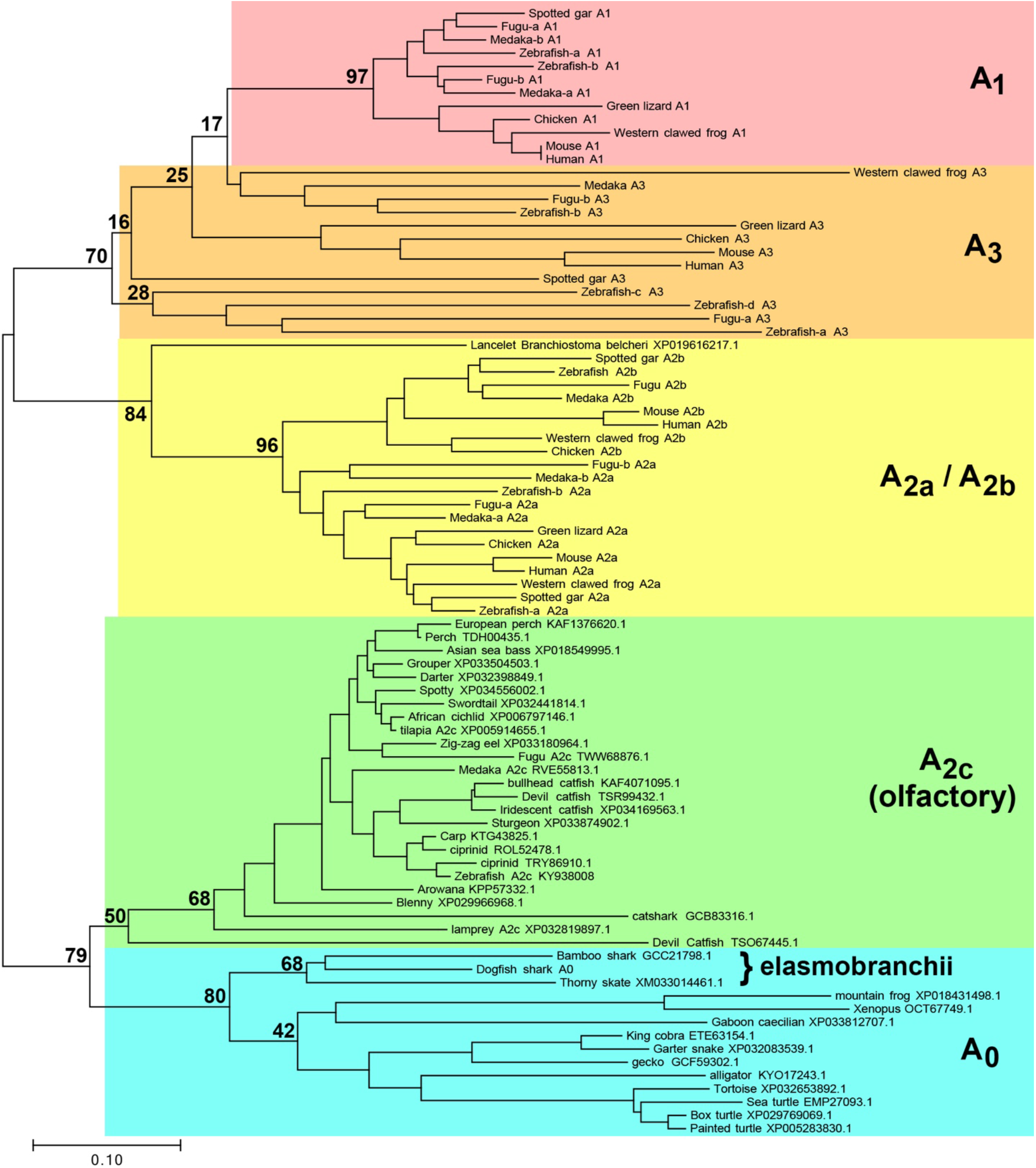
Phylogenetic relationships of ADOR taxa. Evolutionary history is inferred using the Neighbor-Joining method (35). The optimal tree is drawn to scale, with branch lengths in the same units as those of the evolutionary distances used to infer the tree. Distances were calculated using the Poisson correction method (36) and are in the units of the number of amino acids substitutions per site. A_1_, A_2a_, A_2b_, A_3_ and olfactory (A_2c_) ADORs shown in Wakisaka, et al. 2017 are shown using the same names as in the paper. The first characterized A2c (Zebrafish, KY938008) by Wakisaka et al. is also shown in the figure. Additional putative ADORs found in this work are shown by species and Accession. The percentage of replicate trees in which the associated taxa clustered together in the bootstrap test using 500 replicates (Felsenstein 1985) are shown at selected branchpoints. Analyses were conducted using MEGA7 (Kumar, et al, 2016) using only full-length sequences for all receptors.

### *Xenopus laevis* expression system

Adult female *Xenopus laevis* were obtained from Nasco and Xenopus 1, and housed in the animal care facilities at Yale University. Surgeries to harvest oocytes were performed 4-7 days prior to experiments according to a protocol approved by the Yale Animal Care Committee. Injection pipets pulled on a Sachs-Flaming Micropipette puller Model PC-84, using Drummond 210/310/510 glass pipettes were beveled to a tip diameter of 10-15 μm. The pipettes were back-filled with mineral oil and attached to a Drummond microinjector, in preparation for cRNA aspiration and oocyte injection. Oocytes were injected with 50 nl (~15 ng) of cRNA encoding shark A_0_- or human A_1_, A_2a_, A_2b_, or A_3_ receptors and 5-10 ng of human CFTR, and subsequently incubated for 3-5 days at 18-20 °C before experiments.

Oocytes were prepared for two electrode voltage clamping by inserting two 0.6-0.9 MΩ, 3 M KCl filled Ag electrodes, pulled on a Sachs-Flaming micro-pipet puller model PC-84, using borosilicated glass obtained from Sutter Instrument (BF120-69-10). Oocytes were perifused with frog Ringer’s, FR, (98 mM NaCl, 2 mM KCl, 1.8 mM CaCl_2_, 1 mM MgCl_2_ and 5 mM HEPES. Adjusted to pH 7.4), which was also used in the preparation of solutions for the pharmacological agents tested. Oocytes were voltage clamped using the Dagan TEV-200 amplifier along with the Dagan LM-12 Lab Interface, on a PC computer running pClamp Version 5.5 software. The Soltec 1242 plotter was used to acquire the data. Oocytes were clamped at −60 mV and the current was measured as a function of time. Moreover, frequent voltage pulses (−120mV to +60mV) were performed to determine the current-voltage (I-V) relationship at any given timepoint. Data were corrected for any capacitive current incurred due to the voltage pulses. Inconclusive data were obtained with the human A_2a_ and A_3_ receptors, which presumably do not couple efficiently to frog G proteins or the proper adenylyl cyclase(s) for CFTR activation, and were omitted from the analysis.

Data analysis was performed using Microsoft Excel. Some of the data were analyzed using an Excel macro provided by the lab of Dr. David Dawson, which allowed the subtraction of the capacitive current from the current voltage relationship, incurred during the voltage pulses. Statistical analysis was limited to standard deviation performed on the concentration response of the shark A_0_ receptor.

### Homology model of the shark A_0_

At the time of this writing there were 48 high-resolution structures of adenosine receptors in the Protein Data Bank (PDB) as determined by either x-ray crystallography or cryo-electron microscopy. All structures are human in origin and only two of the four known subtypes were represented, A_2a_ (n=45) and A_1_ (n=3). We chose a small subset for consideration as templates for homology modeling of the shark A_0_ receptor: A_2a_ with adenosine bound (PDB code 2ydv) (19), A_1_ with adenosine (6d9h) (20), A_2a_ with N-ethyl-5’-carboxamido adenosine, NECA (6gdg) (21), and A_2a_ with CGS21680 (4ug2) (22). Primary amino acid sequence alignments between shark A_0_ sequence and the selected subset were performed using NCBI Cobalt (23). Amino acid sequences present in the human receptors but not present in shark A_0_ were removed (see full alignment between shark A_0_ and human A_2a_ in supplementary fig. S6, https://www.biorxiv.org/content/biorxiv/early/2020/11/02/2020.11.01.363762/DC1/embed/media-1.pdf?download=true). The human A_2a_ receptor appeared to be a sufficiently good template for A_0_ modeling as it shares 45% identical amino acid sequence with A_0_ and exhibited an NCBI Cobalt 3-bit amino acid chemical similarity of 81% after removing disordered sequences from the A_2a_ structure, and after removing sequences present in either receptor but not the other (supplementary fig S7, https://www.biorxiv.org/content/biorxiv/early/2020/11/02/2020.11.01.363762/DC1/embed/media-1.pdf?download=true). Using the A_2a_ x-ray crystal structure template (PDB code 2ydv), A_2a_ insertion sequences were removed and mutagenesis to A_0_ was performed with scripts written in perl and PyMOL version 2.0.6 (Schrödinger, LLC). Two short amino acid sequences of shark A_0_ that are not present in the human receptors (i.e. A_0_-Arg95 and A_0_-Asn228-Ser230), were inserted into the 3-dimensional model using Coot version 0.8.9 (24) and model minimization/regularization. Refinements using phenix (25) of A_0_ against A_2a_ dataset (PDB code 2ydv) were performed by replacing the human receptor with the shark starting model. Three rounds of rigid body, XYZ-reciprocal space and B-factor refinement were performed including secondary-structure restrains and Ramachandran/rotamer optimization, all with resolution cutoff set to 4.0 Å resolution. Parameterization of the R-PIA ligand was accomplished using phenix (25) starting with the Simplified Molecular-Input Line-Entry System (SMILES) string for the ligand (26): C[C@H](Cc1ccccc1)Nc2ncnc3n(cnc23)[C@@H]4O[C@H](CO)[C@@H](O)[C@H]4O The position of the lipid bilayer was predicted based on the location of conserved aromatic amino acids known to interact with lipid head groups spaced ~35 Å apart (27).

## Results

### Identification of shark A_0_ ADOR

Since the *Squalus acanthias* genome is not yet available and there is some uncertainty associated with the search for *bona fide* protein-coding sequences from genomic data, we sought to clone shark adenosine receptors using reverse-transcription procedures from mRNA/cDNA prepared from shark tissues. PCR of shark rectal gland cDNA using degenerate primers to conserved regions of transmembrane domains 5 and 7 of previously cloned ADORs yielded a single 352 bp product (see supplementary fig. S1). The translated amino acid sequence of this fragment (P-SH12) had highest homology to ADORs that was only 39-45% identical to the human receptor sub-types (A_1_, A_2a_, A_2b_ and A_3_). Northern blot and quantitative RT-PCR (qRT-PCR) revealed strongest expression of the putative shark ADOR in the rectal gland and stomach with weak levels in other tissues including the brain (supplementary figs. S2 and S3). To control for the quality of cDNA template used in the qRT-PCR experiments, shark CFTR was amplified from the same cDNA templates prepared from the various shark tissues. Results showed abundant CFTR expression in rectal gland, intestine, brain and testis (supplementary fig. S4).

Screening of a shark rectal gland cDNA library using 32P-labeled P-SH12 yielded more nucleotide sequence from a single putative ADOR and allowed the design of primers for Rapid Amplification of cDNA Ends-PCR (RACE-PCR). Extensive rounds of both cDNA library screening and RACE-PCR using total RNA from shark rectal gland revealed only a single putative ADOR entity which we designate here as the shark A_0_ receptor (for details see supplementary material).

Hydropathy analysis of the putative shark A_0_ ADOR amino acid sequence revealed seven predicted transmembrane regions and a predicted extracellular orientation of the N-terminus that was consistent with all known G protein-coupled receptors (see supplementary fig. S5). Phylogenetic analysis of known- and putative- ADORs revealed an ancestral branch that included dogfish shark A_0_ and other putative ADORs from non-mammalian species that are distinct from the recently discovered olfactory ADORs (Wakisaka et al. 2017; figure 1). Ancestral olfactory- and non-olfactory receptors could be distinguished from the four specialized mammalian subtypes (i.e. A_1_, A_2a_, A_2b_ and A_3_) as having an abbreviated second extracellular loop (ECL2) at 9 amino acid (aa) residues (supplementary fig. S8, https://www.biorxiv.org/content/biorxiv/early/2020/11/02/2020.11.01.363762/DC1/embed/media-1.pdf?download=true). ECL2 of A_1_ is elongated by 4-17 aa (n=42 receptor species examined), ECL2 of A_2a_ is elongated by 8-14 aa (n=22), ECL2 of A_2b_ is elongated by 10-19 aa (n=28) and ECL2 of A_3_ is elongated by 1-7 aa (n=18, see supplementary fig. S9, https://www.biorxiv.org/content/biorxiv/early/2020/11/02/2020.11.01.363762/DC1/embed/media-1.pdf?download=true). Assuming mutagenesis rates of all adenosine receptors are similar amongst species, we interpret the phylogram to indicate that the elasmobranch A_0_ ADOR belongs to an evolutionary ancestral branch of the specialized adenosine receptor subtypes. We next surmised that shark A_0_ is responsive to adenosine analogs with potentially unique properties compared to known receptor subtypes, and we sought to test this hypothesis using functional and pharmacological characterization in *Xenopus laevis* oocytes.

### CFTR expression

Expression studies were conducted in oocytes co-injected with human CFTR and shark A_0_ cRNA, and current-voltage (I-V) relationships were determined by imposing voltage pulses (−120 mV to +60 mV) and measuring the clamping current. Expression of CFTR was determined by the use of forskolin, a well-known activator of adenylyl cyclase that increases intracellular cAMP levels and activates CFTR by phosphorylation of the R-domain by a cAMP dependent protein kinase A (Dulhanty and Riordan 1994). Figure 2 illustrates the I-V relationships of a single representative experiment on an oocyte 3 days after co-injection. Addition of 5 μM forskolin produced an increase in conductance (increase in the slope of the I-V relationship) from the basal level, an increase in the magnitude of the clamping current at −60 mV from −76 nA to −563 nA, and a rightward shift in the reversal potential to −29.4 ± 0.75 mV (n=3 I-V ramps ± SD) from a basal value of −38.0 ± 3.8 mV (n=5 ramps ± SD). This negative, non-rectifying current with a reversal potential close to that of Cl^−^ (Cl^−^_rev_ = ~ −30 mV) has the electrophysiological characteristics of a CFTR Cl^−^ current. Furthermore, this response was sensitive to 300 μM glibenclamide, an inhibitor of the CFTR Cl^−^ channel (28) (n=1, data not shown). The response to forskolin was also inhibited by the PKA antagonist H-89 (25 μM, n=3) in oocytes injected with human CFTR cRNA (data not shown), suggesting that CFTR is activated by a cAMP dependent protein kinase (PKA) in this system. To confirm, we measured total cAMP in oocytes expressing the shark A_0_ receptor and observed an increase in total cAMP that paralleled the order of agonist potency we observed in electrophysiology experiments (see supplementary fig. S10). The negative, non-rectifying CFTR current was absent in un-injected- and water injected- oocytes (n=4) under identical conditions, and were reproducible both in the same oocyte and in all oocytes injected with human CFTR cRNA. These data established expression of CFTR in our system.

**Figure 2.**
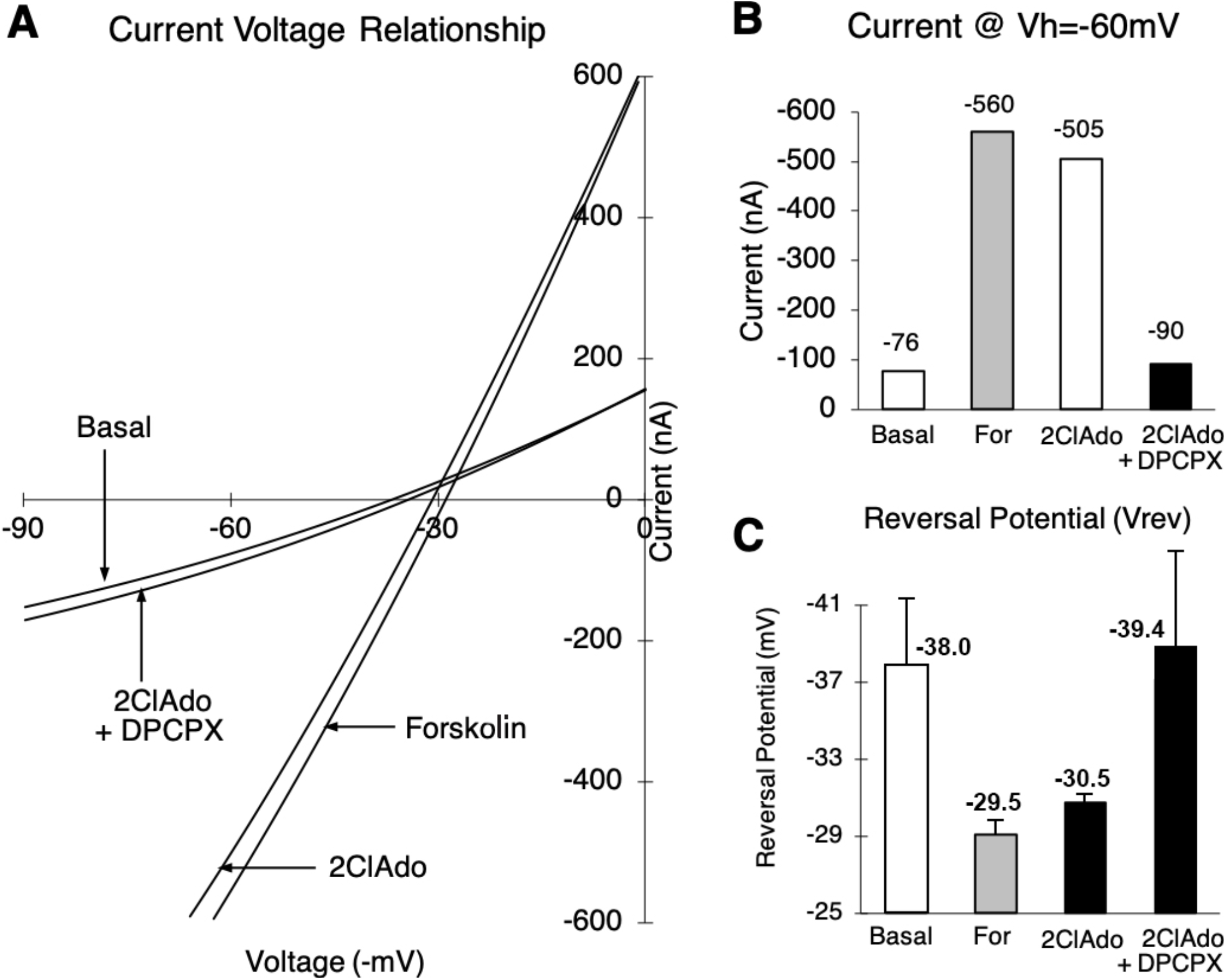
Current-Voltage (I-V) relationship illustrating the effects of forskolin, 2-chloroadenosine (a deaminase-resistant form of adenosine) and 2-chloroadenosine & DPCPX (a known adenosine receptor antagonist) in a representative oocyte co-expressing the shark A_0_ receptor and human CFTR Cl-channel. Panel A: selected single I-V plots for basal, forskolin, 2ClAdo and 2ClAdo+DPCPX. Panel B: Corresponding total oocyte current at −60 mV clamping voltage for the plots in panel A. Panel C: Reversal potentials (mean ± SD) were obtained for basal (−38.0 ± 3.8, n=5), forskolin (“For”, −29.5 ± 0.74, n=3), 2ClAdo (−30.5 ± 0.28, n=4) and 2ClAdo+DPCPX (−39.4 ± 4.8, n=3).

Addition of 10 μM 2-chloroadenosine in the representative oocyte (figure 2) also produced an increase in total current by a similar non-rectifying negative current of the magnitude ~ −500 nA and a reversal potential of approximately −30.5 ± 0.28 mV (n=3 ramps ± SD). This response was reversibly inhibited by 10 μM DPCPX, an adenosine receptor antagonist, where the clamping current at −60 mV was ~ −90 nA with a reversal potential of −39.4 ± 4.8 mV (n=3 ramps ± SD), both of which are reminiscent of the basal state (figure 2). This response suggested that the expression and stimulation of the shark A_0_ receptor is associated with an increase in Cl^−^ current which has characteristics consistent with CFTR C1-currents. Our results also established a coupling of the shark A_0_ adenosine receptor and human CFTR in the *Xenopus* oocyte expression system.

### Pharmacological characterization of shark A_0_

Determination of the pharmacological profiles of adenosine receptor agonists and antagonists in the oocyte necessitated the development of protocols in the *Xenopus laevis* oocyte expression system. We first compared three agonists in a series of identical experiments to provide quantitative data for statistical analysis. We then developed a second protocol to determine a relative response to a larger number of agonists and antagonists to provide a more comprehensive profile for comparison with other receptor profiles in the literature.

#### Quantitative concentration response to agonists

Based on preliminary data, NECA, 2-chloroadenosine (2ClAdo) and R-PIA were selected to determine potency order in a quantitative manner. The concentration response to each agonist was measured as in the representative experiment shown in figure 3. Oocytes were first stimulated with sub-maximal 5 μM forskolin, and subsequently brought to a basal state with frog Ringer’s (FR). The forskolin concentration was chosen to avoid desensitization of the adenylyl cyclase/CFTR pathway since each oocyte was stimulated multiple times by forskolin and adenosine analogs. Next, the oocytes were exposed to increasing concentrations of NECA (0.01 μM, 0.1 μM, 1 μM & 10 μM), each incremental concentration of the agonist was added once the response to the prior concentration had peaked (n=3). This sequence was followed by an identical protocol for 2-chloroadenosine (n=3) and R-PIA (n=6). Variation in the sequence of the agonists tested gave similar results.

**Figure 3.**
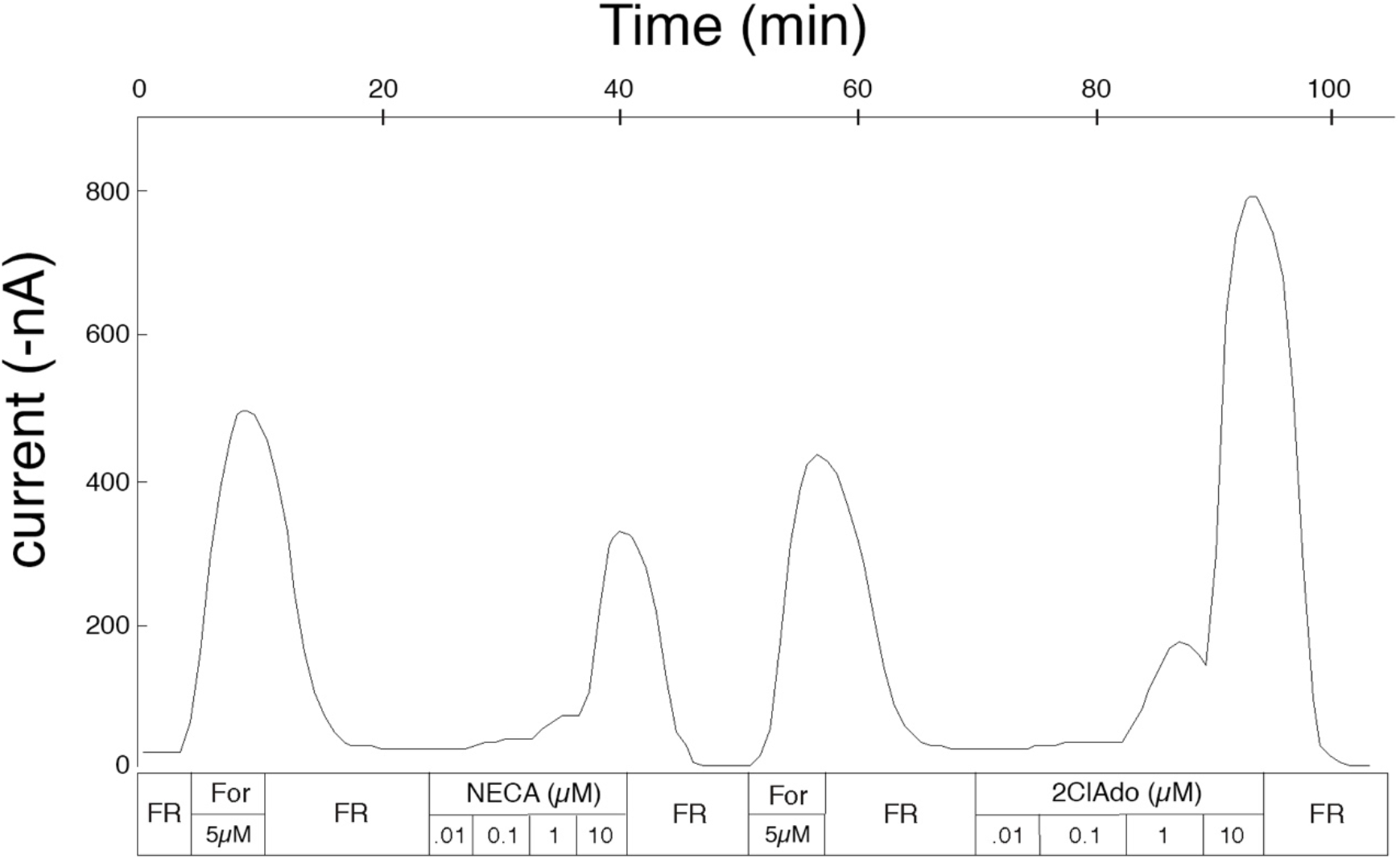
Representative tracing for the determination of the concentration response profile for the shark A_0_ adenosine receptor. A total of four oocytes were used for the determination of the concentration response reported in the main text. Three oocytes, including results from one representative shown in the figure, were exposed to multiple sequences of incremental concentrations of NECA, 2-chloroadenosine and R-PIA (not shown in the figure), while the fourth was used for the study of R-PIA exclusively. Following any stimulation of the oocytes (with For or adenosine analogs), oocytes were returned to basal levels by perifusion with FR.

The initial 5μM forskolin response was used as an internal control (comparison of the agonist response to forskolin) in these studies because of the variability of the expression of CFTR secondary to imprecision in injection of RNA, inter-oocyte variability in CFTR expression and inter-frog and intra-frog variability in the quality of the oocytes due to environmental factors. Data were normalized to the response to 5μM forskolin preceding each concentration response.

Results from the concentration response experiments are summarized in figure 4. The peak normalized stimulation of the control oocytes (injected with human CFTR cRNA only) at 10 μM of NECA, 2-chloroadenosine and R-PIA are −0.01%, 0.04% and 0.03% respectively (normalized to 5 μM forskolin responses, n=2 each, with mean±SD of 191±26 nA, 231±27 nA and 295±96 nA, respectively), showing very poor coupling of any endogenous oocyte adenosine receptor to human CFTR. The responses of oocytes injected with human CFTR and shark A_0_ cRNA, at concentrations of 0.01 μM of NECA, 2-chloroadenosine and R-PIA, and 0.1 μM NECA and 2-chloroadenosine were not significantly different from un-injected controls (p>0.05). The response to R-PIA at 0.1 μM (55.6%) was greater than NECA and 2-chloroadenosine (2.7% and 4.2% respectively). At an agonist concentration of 1 μM the differences between the three agonists NECA, 2-chloroadenosine and R-PIA (13.0%, 35.1% and 196%, respectively) were appreciable and significant (p<0.05). The response at 10 μM (66.5%, 151% and 210% respectively) were markedly different (p<0.05). The probability values for determining a significant main effect for each of these observations were obtained using an ANOVA. These data established the potency order of R-PIA>2-chloro-adenosine>NECA for the shark A_0_ adenosine receptor in *Xenopus laevis* oocytes.

**Figure 4.**
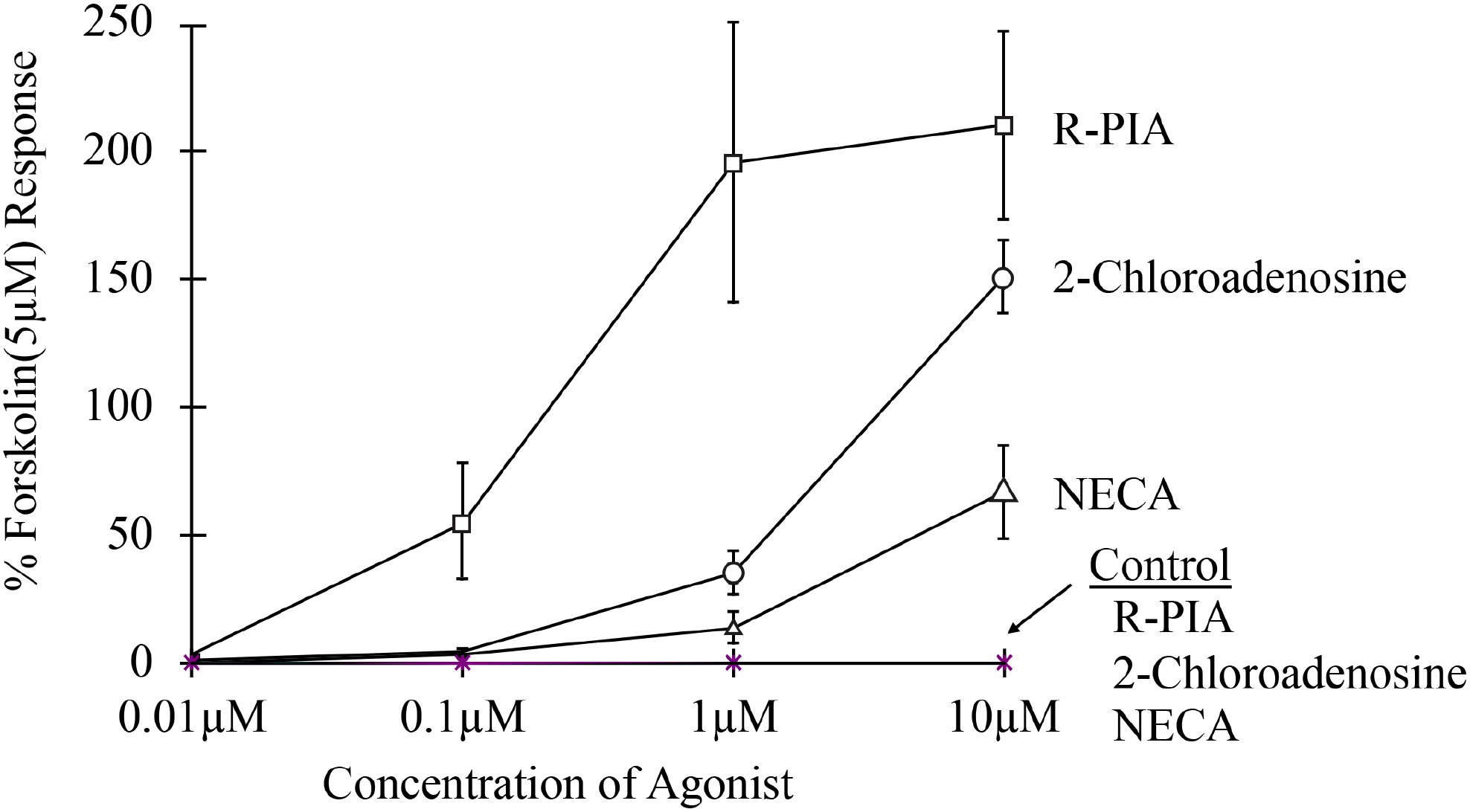
Concentration response of the shark A_0_ adenosine receptor in both A_0_- and CFTR-co-injected oocytes as well as CFTR only-injected oocytes (labeled “Control” in the figure). The controls were conducted in three oocytes, with each agonist tested in each of the three oocytes (see asterisks (*) on the x-axis). The experimental group was tested in four oocytes (NECA, n=3; 2-chloroadenosine, n=3; R-PIA, n=6; and are highlighted in open squares, open triangles and open triangles, respectively). The y-axis is expressed in terms of the percent response to 5 μM forskolin in total oocyte current (nA).

#### Qualitative profile for agonists and antagonists

With the above profile established for the three agonists, a more extensive profile was ascertained for 8 agonists (NECA, DPMA, CV1808, 2-chloroadenosine, CPA,S-PIA, CGS21680, R-PIA) and 5 antagonists (CSC, 8PT, DPCPX, CGS15943, PD115199) to allow comparison with profiles established for the presently cloned adenosine receptors. The analysis was carried out at an agonist concentration of 1 μM and an antagonist concentration of 10 μM to optimize the analysis.

The protocol required oocyte perifusion with one agonist at 1 μM and substitution with a second after stabilization of the signal. The responses were judged as either an increase, a decrease or equivocal. The data were compiled into the agonist profile for the shark A_0_ receptor of R-PIA>S-PIA>CGS21680>CPA>2-chloroadenosine>CV1808=DPMA>NECA (n=7). Varying the order of agonists produced highly consistent results, in that the profiles conducted in order of increasing agonist potency were usually no different from profiles conducted in order of decreasing agonist potency.

The antagonist order of potency was carried out using a similar protocol. Oocytes were perifused with 1 μM R-PIA to generate a stable stimulated state, followed by perifusion with 1 μM R-PIA and 10 μM of the various antagonists. The antagonist profile obtained for the shark A_0_ receptor was DPCPX>PD115199>8PT>CSC>CGS15943 (n=5). The relative potency order of agonists- and antagonists- were determined qualitatively in 10 different oocytes including 186 pairwise comparisons (please see supplementary tables S3, https://www.biorxiv.org/content/biorxiv/early/2020/11/02/2020.11.01.363762/DC1/embed/media-1.pdf?download=true), and 91% (n=169) of all pairwise observations agree with our final conclusion of potency order.

### Protocol validation using human ADORs

The human A_1_, A_2a_, A_2b_ and A_3_ receptors, obtained from the Linden lab at the University of Virginia, were also examined in our oocyte system using the qualitative protocol to provide validation for this form of analysis. Although four human ADORs were expressed in the oocyte, reproducible profiles were obtained for two receptors, the human A_1_ (agonist profile, n=3) and human A_2b_ (agonist profile, n=4, antagonist profile, n=3) (table 1). The human A_1_ and A_2b_ receptor agonist profiles determined in oocytes are in excellent agreement with binding studies and functional studies (see supplementary table S4, https://www.biorxiv.org/content/biorxiv/early/2020/11/02/2020.11.01.363762/DC1/embed/media-1.pdf?download=true). However, the A_2b_ antagonist profile shows an apparent discrepancy in the potency order of CGS15943 and 8PT. This difference may be attributable to a cross species comparison (rat vs human). Taken together, this data provides strong support for the validity of our protocols used in pharmacological characterization of human- and shark- ADORs.

### Homology model of shark A_0_

Given the reasonable sequence homology of shark A_0_ between ADORs that have been well-characterized and for which structures are known (supplementary fig. S11, https://www.biorxiv.org/content/biorxiv/early/2020/11/02/2020.11.01.363762/DC1/embed/media-1.pdf?download=true), we conducted exploratory refinements of the A_0_ starting homology model against the deposited structure factors of the human A_2a_ x-ray crystal structure (PDB 2ydv), replacing the coordinates of A_2a_ with A_0_. Refinement was restricted to low-resolution (4.0 Å) to effectively down-weight the contribution of mis-matched side chain sequences between the two species while providing a data-driven constraint for backbone atoms and secondary structure. The resulting model exhibited good protein Ramachandran- and side chain rotameric geometry (supplementary table S5, https://www.biorxiv.org/content/biorxiv/early/2020/11/02/2020.11.01.363762/DC1/embed/media-1.pdf?download=true) and had convincing secondary structure for the transmembrane helices and extramembrane segments (figure 5). Both sequence alignments and the shark A_0_ homology model revealed an abbreviated extracellular loop 2 (ECL2) in ancestral receptors compared to the more specialized subtypes. While it is not yet known what, if any, functional consequence arises from different lengths of the ECL2 specifically, more than 30 ancestral receptors in the database have a minimized ECL2 (supplementary fig. S8). Thus, it would seem that ECL2 of ancestral receptors is at the minimum length necessary to maintain the structure and/or function of the adenosine receptor architecture.

**Figure 5.**
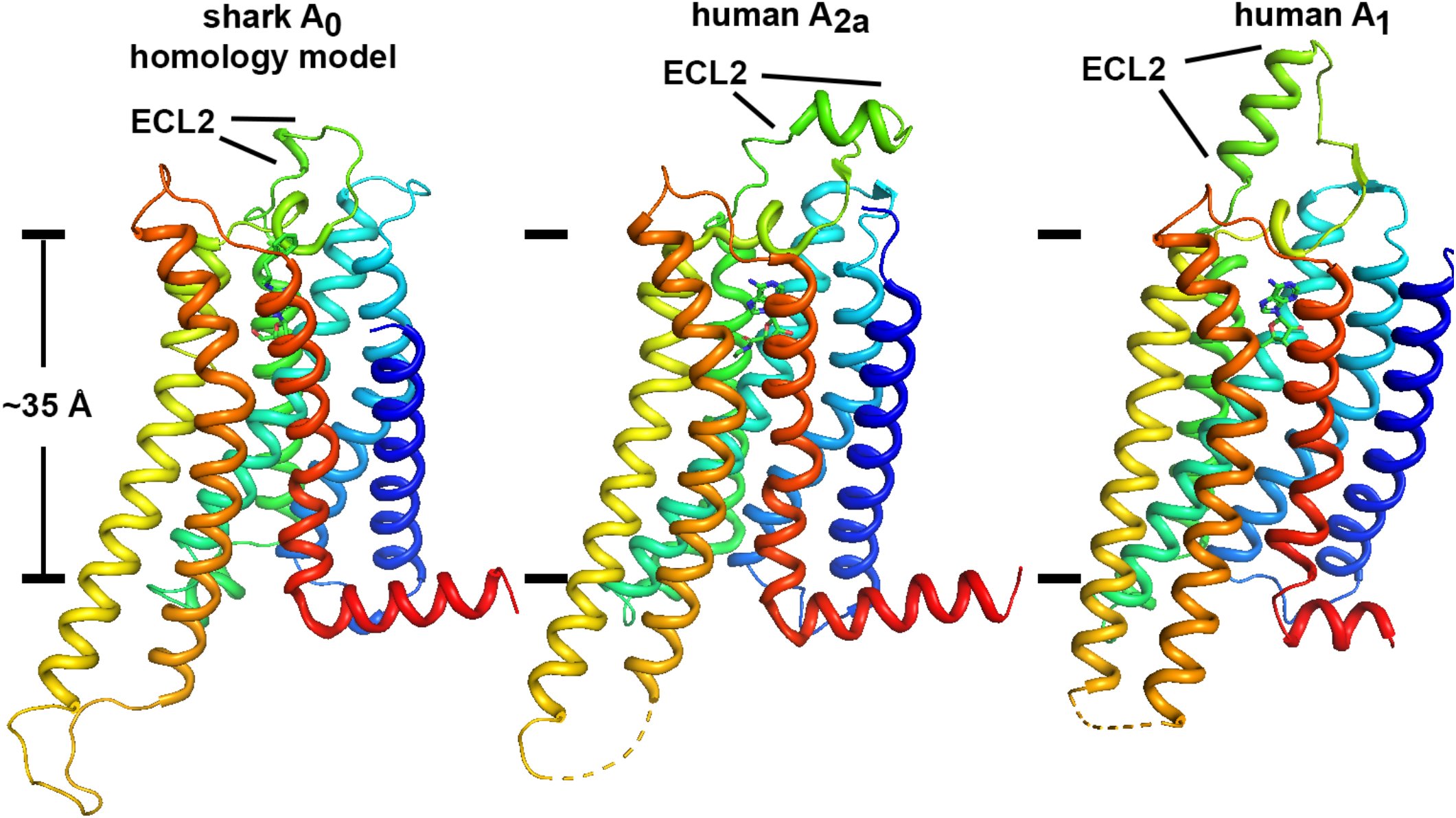
Overall view of the shark A_0_ homology model compared to human A_2a_ and A_1_ receptors (PDB codes 2ydv and 6d9h). The approximate position of the hydrophobic stretch of the lipid bilayer is depicted with lines separated by ~ 35 Å using conserved aromatic side chains as fiducials (27). A striking feature of shark A_0_ compared to the more specialized ADORs is an abbreviated extracellular loop 2 (ECL2, labeled) and the relatively low profile that the A_0_ receptor presents to the extracellular milieu. Multiple sequence alignment of all known- and putative receptor orthologs (supplementary figs. S9 and S10) reveals that an abbreviated ECL2 appears to be unique feature of ancestral ADORs.

### Conserved Ligand-Receptor Interactions

To explore the extent to which our homology model recapitulates known fundamental interactions between ligand and receptor, we separately modeled adenosine (ADO), 2-chloroadenosine and the most potent ligand in this work, R-phenyl-isopropyl adenosine (R-PIA), into the A_0_ binding pocket, conducted refinements and inspected the receptor-ligand interactions. While we recognize there will be error in the process of homology modeling A_0_ in the absence of an actual structure, potential fundamental interactions between A_0_ and ligands are well conserved in the refined A_0_ model compared to human ADOR structures (figure 6 and supplementary table S6, https://www.biorxiv.org/content/biorxiv/early/2020/11/02/2020.11.01.363762/DC1/embed/media-1.pdf?download=true).

**Figure 6.**
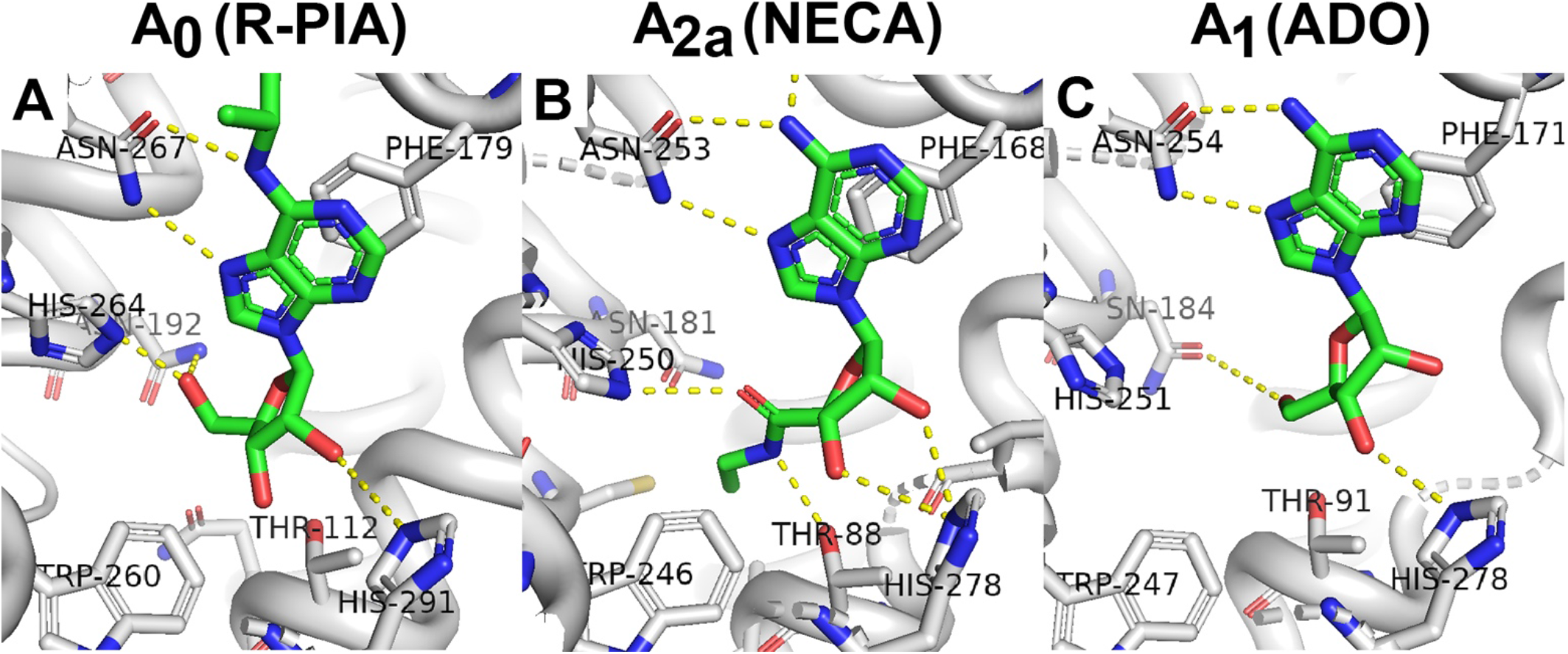
Binding pockets for adenosine and adenosine agonists. Panel A shows a potential binding mode of R-phenyl-isopropyl adenosine (R-PIA) as it appears in our refined homology model of the shark A_0_ receptor. The purpose is not to highlight precise interactions, bonds or geometry, but to indicate that the major known adenosine-binding ligands are conserved in the shark A_0_ ancestor receptor and are well positioned to potentially interact with ligands. For comparison, the location of N-ethyl-carboxyl adenosine (NECA) binding in the 2.6 Å x-ray structure of human A_2a_ receptor (PDB code 2ydv) is shown in panel B. Panel C shows the location of adenosine (ADO) binding in the 3.6 Å cryo-EM structure of the human A_1_ receptor (PDB code 6d9h).

Among the most important known interactions between receptor and ligand occur between ADOR and the purine pharmacophore of adenosine- and adenosine analogs. The structures of human A_2a_- and A_1_ reveal that an aromatic residue (F168 and F171, respectively), protrudes down into the ligand binding pocket and contribute an intimate π-π stacking interaction with the purine ring of adenosine (29). Our refined homology model of shark preserves this interaction convincingly well, with the corresponding A_0_ aromatic residue being F179 (figure 6). A second critical interaction revealed by the structures of A_2a_- and A_1_ is a strong polar interaction, most likely hydrogen bonding, between an asparagine residue (N253 and N254, respectively) and the N7 nitrogen and the amino group of C6 of the purine ring. The corresponding A_0_ asparagine is N267 (figure 6). These two interactions form the fundamental basis for recognizing the adenosine pharmacophore and are invariant in all ADORs we have examined (see multiple sequence alignments in supplementary figs. S12 and S13, https://www.biorxiv.org/content/biorxiv/early/2020/11/02/2020.11.01.363762/DC1/embed/media-1.pdf?download=true). In totality, nine amino acid residues in close proximity to the adenosine analog in our shark A_0_ homology model are highly conserved in all known- and putative ADORs, including the ancestral branch (see also supplementary table S6).

## Discussion

### Co-expression of shark A_0_ and human CFTR

We successfully co-expressed the shark A_0_ adenosine receptor with human CFTR in *Xenopus laevis* oocytes, which, to our knowledge, represents the first heterologous co-expression of any adenosine receptor with CFTR in the literature. Oocytes appear to be an ideal system to characterize shark A_0_, since neither frog ADORs nor frog CFTR appears to be active or coupled. We coupled an adenosine receptor from an elasmobranch shark, a subclass of Chondrichthyes fish with a temporal range dating back to at least the Devonian period (~419-360 million years ago), to human CFTR using the endogenous G-proteins and cellular enzymes of the oocyte. Other investigators have in the past exploited CFTR for co-expression studies, specifically with alpha 2-adrenergic-, beta 2-adrenergic-, delta-opioid-, and the 5HT1A receptors ((30), (31)), however those receptors were mammalian in origin, which would not be viewed nearly as long-lived as receptors identified in the shark. A_0_ function in oocytes suggests that the interaction between receptors and frog G-proteins is maintained despite the ancestral properties and structural differences A_0_ has with respect to more specialized ADORs. This highlights a likely conservation of the interface between G-proteins and receptors with which they interact, across species and through evolution. Our results also emphasize the versatility and value of the *Xenopus laevis* expression system for studies of GPCR molecular evolution.

### A_0_ exhibits unique ADOR pharmacology

We characterized the shark A_0_ receptor in *Xenopus* oocytes using a novel protocol, in contrast to previous studies using binding assays, *in vivo* functional studies and cAMP measurements. To this end it was necessary to couple the shark A_0_ with a reporter target protein, in this case human CFTR, to permit measurement of expression by electrophysiology. The data from these studies revealed a unique profile for the shark A_0_ receptor. Table 2 summarizes the comparison of the A_0_ receptor to other adenosine receptors with respect to agonist potency.

The shark A_0_ differs: 1) from A_1_ in its potency order of 2ClAdo and NECA, 2) from A_2a_ and A_2b_ which have the reverse potency order and 3) from A_3_ in which the least potent shark A_0_ agonist, NECA, and the most potent shark A_0_ agonist, R-PIA, have equal potencies. Therefore, the shark A_0_ appears to have a unique agonist profile judging from the potency order of R-PIA > 2ClAdo > NECA.

Subsequent qualitative studies further established the unique nature of the shark A_0_. Table 3 compares the pharmacological profile for the shark A_0_ to mammalian A_1_, A_2a_, A_2b_ and A_3_ adenosine receptor subtypes. Analysis reveals the A_1_ specific agonist R-PIA is the most potent in the shark, however the A_1_ selective agonist CPA is of intermediate potency. While S-PIA and CGS21680 are more potent in the shark A_0_ than in the A_1_ receptors. All of the above are inconsistent with an A_1_-like receptor profile for A_0_.

The shark A_0_ receptor is dissimilar to A_2a_ or A_2b_ with respect to its low potency to the A_2_-preferring agonist NECA. Furthermore, response of A_0_ to S-PIA was unexpectedly high considering that S-PIA is not relatively potent at A_1_, A_2a_ or A_2b_ receptors. Interestingly, the A_2a_ preferring agonist CGS21680 has relatively high potency in the shark A_0_ profile, while CV1808, also A_2a_ preferring, has a much lower potency in the profile. The shark A_0_ profile bears little resemblance to that of A_2b_ receptors especially with respect to the position of CGS21680, NECA, R-PIA and S-PIA. Therefore, the shark A_0_ resembles neither the A_2a_ nor the A_2b_ adenosine receptors (table 3). Similarly, it is quite different from the known A_3_ adenosine receptor profiles (table 3), with low potency of NECA (intermediate potency in A_3_), relatively high potency of CGS21680 and 5-PIA (lower in the profile for A_3_) and the large difference in potency of R-PIA and NECA, which are of equal potency in the A_3_ adenosine receptor profiles.

A comparison of the antagonist profiles yields similar results. Although the shark A_0_ antagonist profile is reminiscent of the A_1_ antagonist functional profile (table 3), when compared to binding studies for the rat A_1_ (supplementary table S7, https://www.biorxiv.org/content/biorxiv/early/2020/11/02/2020.11.01.363762/DC1/embed/media-1.pdf?download=true), the low potency of CGS15943 in the shark A_0_ profile is striking. The relative potency of CGS15943 is also incongruent with the rat A_2a_ and rat A_2b_ profiles (supplementary table S8, https://www.biorxiv.org/content/biorxiv/early/2020/11/02/2020.11.01.363762/DC1/embed/media-1.pdf?download=true). Moreover, the potency order of DPCPX<PDl15199 in the shark A_0_, is reversed for A_2a_ receptors and A_2b_ receptors. It is clear that shark A_0_ also possess a unique antagonist profile.

In summary the agonist- and antagonist- profiles do not allow the classification of this receptor into any known receptor subtype, providing functional evidence that A_0_ is a unique ADOR.

### A_0_ is an extant ADOR ancestor

Since the initial characterization of specialized adenosine receptor (ADOR) subtypes ~40 years ago, hundreds of ADORs have been identified. Many examples arise from genomic sequencing but have not been characterized structurally or functionally. Phylogenetic analysis shows that examples of specialized ADORs (i.e. A_1_, A_2a_, A_2b_ and A_3_) are apparent throughout the animal kingdom, including fish, amphibians, reptiles, birds and mammals. However, a separate phylogenetic branch emerges from the analysis that contains ADORs in reptiles, amphibians and fish, including the shark A_0_ ADOR we have characterized here. Shark A_0_, and the related members on the branch, appear to be ancestral in nature. We draw this conclusion based on the following: 1) amino acid sequence similarity of A_0_ is roughly equally different from any specific specialized subtype (A_1_, A_2a_, A_2b_ and A_3;_ see supplementary table S9, https://www.biorxiv.org/content/biorxiv/early/2020/11/02/2020.11.01.363762/DC1/embed/media-1.pdf?download=true); 2) the pharmacological profile of A_0_ is significantly different from the profiles of any subtype; 3) A_0_ belongs to a phylogenetic branch that is distinct from the branches formed by the other specialized subtypes; and 4) ADOR members of the ancestor branch contains structural features unique only to themselves, such as a minimal ECL2.

A_1_, A_2a_, A_2b_ and A_3_ appear to be specialized for several reasons: 1) they have distinct and consistent pharmacological profiles amongst their respective sub-branches; 2) they prefer to couple to one G-protein alpha subunit (e.g. inhibitory G_i_α or stimulatory G_s_α on adenylyl cyclase activity), and 3) they are abundant in birds and mammals which are thought to arrive later in evolution. The latter point highlights an interesting observation: specialized ADORs are prevalent in mammals which arrived more recently in evolution time-scales, but the ancestral receptors, including olfactory- and non-olfactory ADORs, appear to have been lost in mammals.

## Conclusion

We have cloned, expressed and characterized an adenosine receptor (A_0_) from the dogfish shark (*Squalus acanthias*) that exhibits novel pharmacology compared to all known subtypes (A_1_, A_2a_, A_2b_ and A_3_). A_0_ and its closest relatives form a distinct ancestral phylogenetic branch from the known subtypes and do not appear have representation in birds or mammals. All ancestral ADORs appear to have distinguishing structural elements, such as a minimally short extracellular loop 2. Shark A_0_ expression is strong in rectal gland where it likely regulates chloride secretion but its expression is weak in brain, allowing us to conclude that is not olfactory in primary function. A non-olfactory ancestral branch of ADORs, including A_0_ from elasmobranchs, amphibians, and reptiles is now apparent from the wealth of genomic sequencing data.

In contrast to the recently identified olfactory ADOR (A2c) found exclusively in fish (13), A_0_ was cloned from the shark rectal gland, a non-olfactory epithelial organ known for diverse GPCR-mediated regulation of chloride secretion via the CFTR chloride channel (32). Shark CFTR is more than 70% identical to human CFTR, and ADOR regulation of CFTR in sharks and mammals is well described (33,34). Taken together, a long-standing evolutionary functional coupling between ADORs and CFTR-mediated chloride secretion in epithelial tissues is evident, and the elasmobranch A_0_ receptor emerges as a non-olfactory extant ancestor of specialized ADOR subtypes that most likely regulates CFTR in the shark.

## Acknowledgements

We wish to thank Drs. Stephanie Halene and Kristina Lübbe for their important contributions toward the manuscript. This work was supported by a Howard Hughes Medical Institute Grant to SB, internal UAB funding to SGA, and by NIH grants R01 DK034208, NIEHS P30 ES003828 and NSF grant DBI-0453391 to JNF.

## Data Availability

We agree to deposit the shark adenosine receptor nucleotide sequence to public repositories (NCBI) or make it available upon request.

