## Supplementary Document for "A non-olfactory shark adenosine receptor activates CFTR with unique pharmacology and structural features"

*Preparation of Total RNA from Rectal Gland Tissue*. Total RNA was extracted from 1 g of flash frozen shark rectal gland tissue. Frozen tissue was ground by mortar and pestle and homogenized in a motor-driven homogenizer (Tekmar) in 10 ml denaturing solution (4 M guanidium thiocyanate, 25 mM sodium citrate, pH 7.0, 0.5% sarcosyl, 0.1 M 2-mercaptoethanol). DNA and proteins were removed by adding 0.1 volume 2 M sodium acetate, pH 4.0, 1 volume buffer saturated phenol, pH 8.0, and 0.2 volume chloroform/isoamyl alcohol (24:1) and centrifuging at 10,000 x g for 15 min at 4 ˚C. Polysaccharides were removed by resuspending the pellet in 4 M LiCl and centrifuging at 3,000 x g for 10 min. The RNA pellet was re-dissolved in TE buffer (10 mM Tris-HCl, pH 7.5, 1mM EDTA, pH 8.0) with 0.5% SDS. An equal volume of chloroform/isoamyl alcohol (24:1) was added and the aqueous layer was collected after centrifugation at 3,000 x g for 10 min. The RNA was precipitated with 2 M sodium acetate and isopropanol at -20 ˚C for 15 min. The precipitate was centrifuged at 10,000 x g for 15 min at 4 ˚C and the RNA pellet was washed with 75% ethanol. The pellet was dried under vacuum and resuspended in TE buffer.

*Degenerate Oligonucleotide Primer Design and Cloning of Shark Adenosine Receptor*. Total RNA was prepared from shark rectal gland tissue as described above and first strand cDNA was synthesized using AMV reverse transcriptase and oligo dT-primers (Invitrogen). Amino acid sequences from previously cloned adenosine receptors retrieved from GenBank were searched by a computer program (S. Aller) to identify regions of high homology and low codon degeneracy. The forward sense oligo primer, ADOF, (5'-ATG(A/G/T)(A/G/C) ITA(C/T)ATGGT(A/G/C/T)TA) codes for the N-terminus of the fifth transmembrane domain of all previously identified adenosine receptors. The antisense primer, ADOR, (5'-GC(G/T)TAIA(C/T)(A/G/T)A(A/T) (A/G/ C/T)GG(A/G)TT) codes for the C terminus of the seventh transmembrane domain. PCR was performed with ADOF and ADOR primers on rectal gland cDNA using Amplitaq (Perkin Elmer) with denaturing for 1 min at 95˚ C, followed by 35 cycles of 95 ˚ C for 1 min, 40 ˚C for 1 min, and 72 ˚C for 1 min followed by final extension at 72 ˚C for 3 min in a Techne PHC-3 thermocycler. The 352 bp PCR product was purified from a 1% low melt agarose gel using glassmilk separation (Bio 101) and cloned into the pCRII TA cloning vector (Invitrogen). The insert was sequenced manually in both directions by the dideoxy chain termination method Sanger, 1977 #173 using Sequenase II (U.S. Biochemical Corp.).

*cDNA Library Construction and Screening*. One gram of frozen rectal gland tissue was homogenized in guanidium thiocyanate buffer (4 M guanidium thiocyanate, 0.1 M Tris-HCl pH 7.5, 1% 2-mercaptoethanol, 0.5% sodium lauryl sarcosinate). The homogenate was layered onto 5.7 M CsCl, 0.1 M EDTA and ultracentrifuged in a Beckman centrifuge (SW41 swinging rotor) at 32,000 rpm for 24 h. The RNA pellet was washed with 70% ethanol, redissolved in TE buffer, pH 7.6, 0.1% SDS re-precipitated with 3 M sodium acetate, pH 5.2, and ethanol for 1 hr at -20 ˚C, and centrifuged at 12,000 g for 30 min at 4 ˚C. The RNA was washed in ethanol, re-centrifuged, dried, and dissolved in water. Poly(A) RNA was purified from total RNA using magnetic bead separation (PolyATtract, Promega Corp). A cDNA expression library was prepared in λZAP II using both oligo(dT) and random primers (custom library, Stratagene). The average insert size of the library was 2.5 kb using vector specific primers on 20 random pure phage colonies. The titre of the unamplified library was $9.6x10^5$ plaques/ml. The gel-purified PCR fragment was radiolabeled with 5'-[a-$^{32}$P]-dCTP (Amersham) using random primer DNA labeling (Gibco). Unincorporated nucleotides were removed with G-50 Sephadex spin columns (Boehringer Mannheim). Library phage were transferred to Hybond-N nylon membranes (Amersham) in duplicate. Membranes were placed in a denaturing solution (0.5 M NaOH, 1.5 M NaCl), washed in 1.5 M NaCl, 0.5 M Tris-HCl, pH 7.4, 10 mM EDTA, pH 8.0 and soaked in 2x SSC for 3 min. The DNA was fixed to the membranes by baking for 2 h at 80 ˚C. Filters were hybridized at high stringency overnight. Filters were then washed with 1 mM EDTA, 1% SDS, 40 mM $NaHPO_4$, pH 7.2, for 1 min at room temperature followed by two washes of 5 minutes at 65˚C and exposed to X-ray film in autoradiography cassettes with intensifying screens for 2 days. Plaques were subjected to three rounds of purification. Inserts of the purified plaques were each excised with helper phage and transformed into XLOLR E. coli strain (Stratagene). Highly pure plasmid DNA was prepared and the nucleotide sequence of the inserts was obtained by manual sequencing as mentioned above using vector specific primers T3 and T7.

*RACE PCR.* Total RNA was prepared from frozen rectal glands as described above. First strand cDNA was reverse transcribed from total RNA using MMLV reverse transcriptase and an oligo(dT) primer cassette. Second strand cDNA was synthesized using RNase H, E. coli DNA polymerase I and E. coli DNA ligase. Blunt ends of the cDNA were generated with T4 DNA polymerase and ligated to cDNA adaptors at both ends of the cDNA (Marathon cDNA Amplification, Clontech). Two shark adenosine receptor-specific oligos were designed from library clone sequence to yield a 5' and a 3' RACE PCR fragment. "RADO5" (5-GAGAGCGAGTTATGGTGCGTC) was an antisense primer corresponding to shark nucleotides 1486-1506. RADO3 (5-GCTTCCTGATG CTGACCACG) was a sense primer corresponding to shark nucleotides 1181-1201. RACE PCR was performed using a primer that anneals to the Marathon cDNA adaptor sequence and either sense or antisense shark adenosine receptor specific primers ("RADO5" or "RADO3"). Expand High Fidelity Taq Polymerase (Boehringer Mannheim) was used for high fidelity thermocycling. A 50 µl reaction mixture was heated to 94 ˚C for 2 min then subjected to 30 cycles of 94 ˚C for 30 s, 60 ˚C for 45 s and 68 ˚C for 3 min. The resulting 5' and 3' RACE PCR products were directly ligated into the pCRII TA cloning vector (Invitrogen). Competent cells (Invitrogen) were transformed with the ligated plasmid and bacterial colonies containing an insert were chosen via blue/white selection. Insert-containing plasmids were prepared from white colonies. The inserts were confirmed to be shark adenosine receptor fragments by automated DNA sequencing as described below.

5'-RACE-PCR resulted in a single band of 1485 bp that included 322 bp of 5'- untranslated region (5'-UTR), an open reading frame of 1056 bp encoding an entire putative adenosine receptor, and 155 bases of the 3'-UTR (supplementary fig. S1B and S1C). A short 3'-RACE product gave further 3'-UTR sequence. The 5'-RACE PCR product was subcloned into *E. coli* and 10 positive colonies were bidirectionally sequenced. Six of 10 plasmids contained identical 370 bp of 5'-UTR, an open reading frame of 1056 bp and a 3'-UTR of 107 bp. Three plasmids had deletions of 20 bases and one was missing 40 bases in the initial portion of the 5'-UTR. All 10 clones had identical sequence in the open reading frame and 3'-UTR. A second screening of the rectal gland cDNA library yielded two additional positive clones, one of which (L-SH5) contained the full open reading frame (1056 bp) encoding the receptor protein (supplementary fig. S1C). The translated amino acid sequence of this library clone was identical to the full length 5'-RACE PCR product.

S*equencing and Analysis.* cDNA clones were bidirectionally sequenced by automated sequencing at the University of Maine (Applied Biosciences Inc.) using Amplitaq FS (Perkin-Elmer) and dye terminators. Samples were subjected to 25 cycles of thermocycling with the following parameters: 96 ˚C for 30 s, 50 ˚C for 15 s and 60 ˚C for 4 min in an MJ Research PTC-100 thermocycler. Hydropathy analysis of the putative shark adenosine receptor amino acid sequence revealed seven clearly predicted transmembrane regions and a predicted outside orientation of the N-terminus that was consistent with all known G protein-coupled receptors (see supplementary figs. S5 and S11 below).

*Northern Blot Analysis*. Total RNA was isolated from flash frozen shark tissues (brain, gill, heart, kidney, liver, muscle, rectal gland, spleen, testis) using the guanidium phenol/chloroform extraction method as described above. 30µg of total RNA from each tissue was denatured and fractionated on a 1.2% agarose/formaldehyde gel in 1 x MOPS buffer. The electrophoresed RNA was transferred to GeneScreen Nylon membranes and fixed by baking at 80˚C for 2 hours. The original 352 bp PCR fragment (P-SH12) obtained from degenerate PCR) was used for hybridizing the membrane. Approximately 50 ng of the PCR product was denatured by heating to 95˚C for 5 min and radiolabeled as described above. The denatured radiolabeled probe was added to the hybridization solution (1% SDS, 1M NaCl, 10% dextran sulfate, 50% formamide) along with 150 µg of denatured salmon sperm DNA. The blot was hybridized at 42˚C for 18 hours. Following hybridization, the membrane was washed stringently in 2x SSC at room temperature for 5 minutes, followed by 2x SSC, 0.5% SDS at 60˚C for 30 min, followed by two washes in 0.1x SSC at room temperature for 30 min each. The membrane was exposed to x-ray film for 7 days.

R*T-PCR*. Total RNA extraction, reverse-transcription and PCR were carried out as above, except that RNA samples were pre-treated with RNAse free DNAse. Control reactions contained either no template RNA or no cDNA. The internal standard was a shark specific histone H3.3 subtype, cloned by our laboratory. Duplicate

PCR reactions were performed on cDNA from each tissue using shark $A_0$ and shark histone H3.3 primers. Results were verified by restriction enzyme digestion of known restriction sites of the $A_0$ receptor.

*Preparation of Adenosine Receptors for Expression Studies.* A sense primer, ADOSTART, (GGGAGAGGAAGACTACGCAAT) that is specific for the shark adenosine receptor nucleotides located 34-13 bp upstream of the start codon was designed as well as a specific antisense primer, ADOSTOP, (CACAGGATCTCTCCCAGCATA) corresponding to nucleotides 2-22 downstream of the stop codon. ADOSTART was phosphorylated and used with ADOSTOP to perform PCR on shark rectal gland cDNA using Expand High Fidelity Taq Polymerase (Boehringer Mannheim). The reaction mixture was heated to 95 ˚C for 1 min before addition of enzyme. 28 cycles were performed at 95 ˚C for 30 s, 55 ˚C for 30 s and 68 ˚C for 4 min. A 1114 bp product was obtained and directly cloned into the pCR3 expression vector (Invitrogen). Competent cells were transformed and plasmid DNA of the positive colonies was prepared. The insert sequence was confirmed as the $A_0$ receptor by automated sequencing (Applied Biosystems Inc.). The plasmid was linearized using Xba I endonuclease followed by phenol/chloroform extraction to inactivate the enzyme and remove impurities. An ethanol precipitation (0.1 volume of 3M sodium acetate, pH 5.0-Promega & 2.5 volumes of 100% ethanol) was carried out to further purify the product. Capped messenger RNA was synthesized from the plasmid DNA template using the mMessage mMachine T7 in vitro transcription kit (Ambion 1344). Purification of the RNA was achieved by isopropyl alcohol precipitation. The RNA product was a 1.35kb transcript. The quality and purity of the final product was assessed by gel electrophoresis.

*Xenopus laevis expression system*. Adult female *Xenopus laevis* were obtained from Nasco and Xenopus 1, and housed in the animal care facilities at Yale University. Surgeries to harvest oocytes were performed 4-7 days prior to experiments according to a protocol approved by the Yale Animal Care Committee. An adult female frog was immersed in cold anesthetic solution (0.16% solution of Tricaine (3-Aminobenzoic acid ethyl ester methane-sulfonate salt adjusted to pH 7.0) for 10 minutes to achieve anesthesia. A few ovarian lobules were removed under sterile conditions from an abdominal incision and collected in collagenase solution (2.5 mg/ml of collagenase A from Boehringer Manheim) in denuding solution (82.5 mM NaCl, 2.5 mM KCl, 1 mM $MgCl_2$, and 5 mM HEPES. Adjust to pH 7.6). The muscle layer was closed with 5.0 chromic gut, and the skin incision with 5.0 nylon. The oocytes were incubated in the collagenase for a 2 to 3 hours (depending on the collagenase activity) at room temperature. Mature stage V & VI oocytes were manually defolliculated using #5 forceps under a dissection microscope and stored in Modified Barth Solution Holding media (MBSH) (88 mM NaCl,1 mM KCl, 2.4 mM $NaHCO_3$, 0.82 mM $MgSO_4$, 0.33 mM $Ca(NO_3)_2{*}4H_2O$, 0.41 mM $CaCl_2{*}2H_2O$, 10mM HEPES and 150 mg/L of Gentamicin sulfate adjusted to pH 7.6).

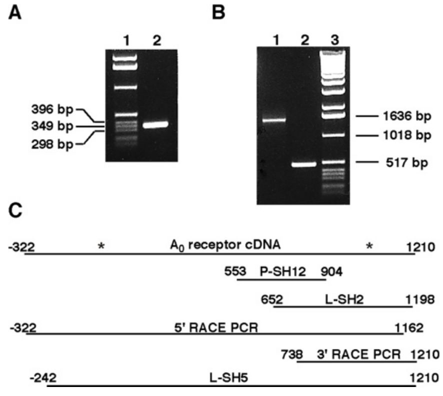

**Figure S1**. Results of PCR and library screening to clone and identify shark adenosine receptors. The original PCR product resulting from degenerate PCR is shown in panel A. RACE (Random Amplification of cDNA Ends) PCR results (5'- and 3'-) are shown in panel B. Panel C shows a schematic of sequences obtained from all rounds of PCR and library screening, with the final published sequence shown for reference at the top of the panel. Fragments L-SH2 and L-SH5 resulted from two independent rounds of cDNA library screening, and amplicons resulting from the RACE-PCR experiments are labeled. All DNA sequences we achieved in our search for shark adenosine receptors revealed only a single putative ADOR in the shark rectal gland.

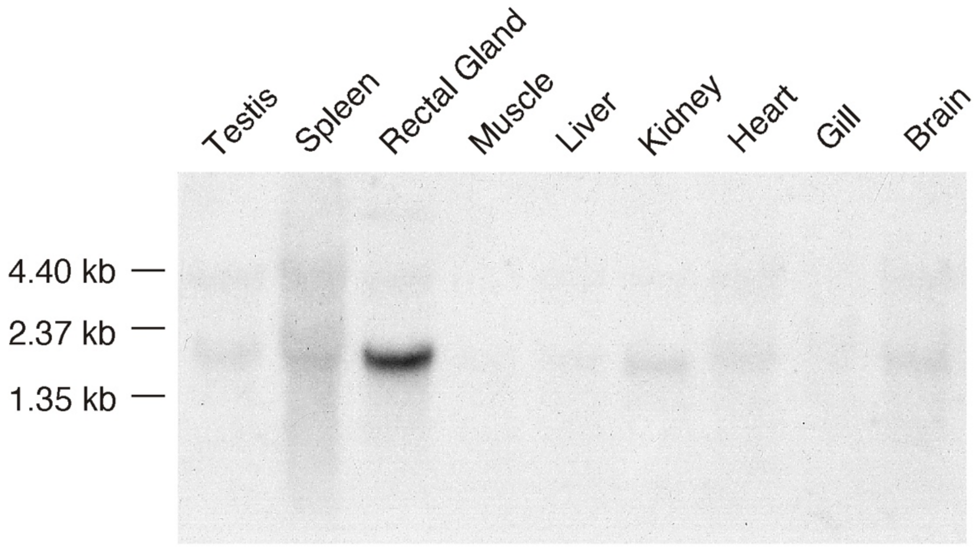

**Figure S2.** Northern blot of total RNA prepared from shark tissues using a $^{32}$P-labeled DNA probe corresponding to the original degenerate PCR product (P-SH12).

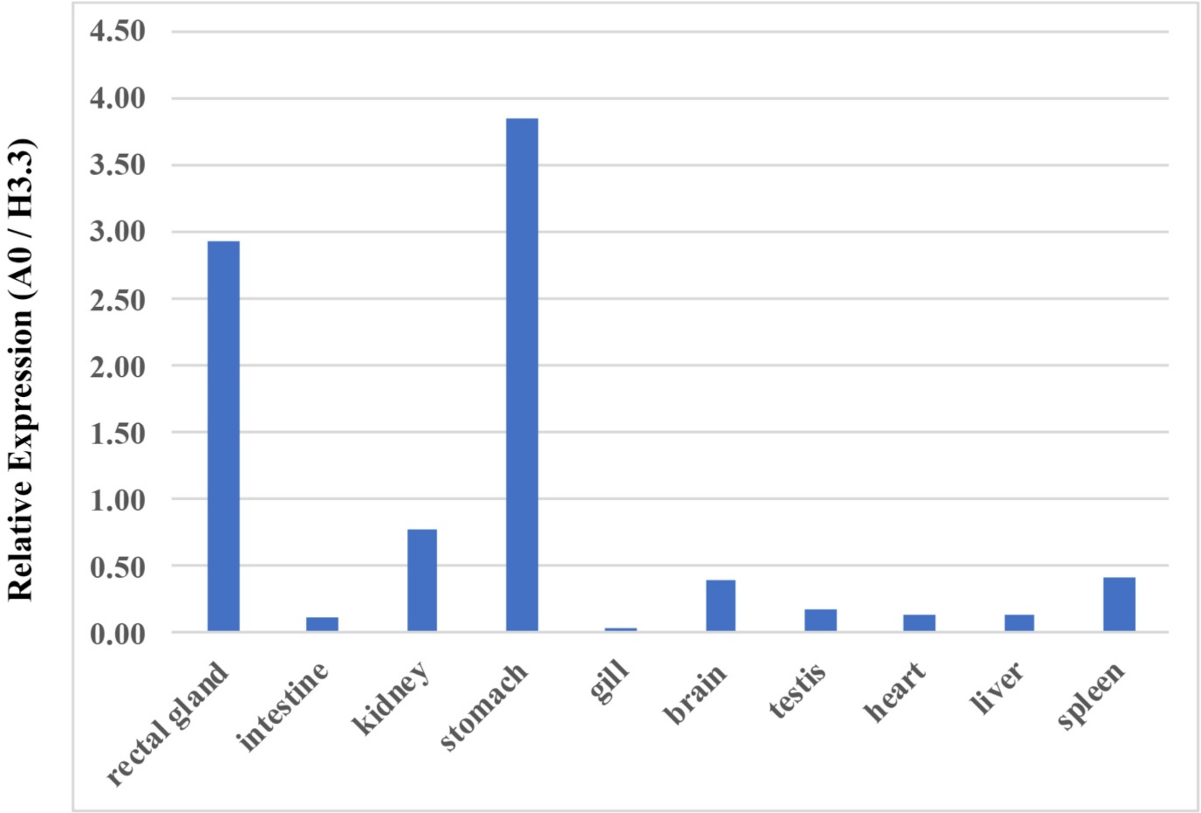

**Figure S3.** Expression of $A_0$ adenosine receptor in selected shark tissues. Complimentary DNA (cDNA) was prepared from the indicated shark tissues and $A_0$ adenosine receptor was amplified by quantitative RT-PCR using $A_0$-specific primers. Shark histone H3.3 was also amplified simultaneously, and densitometry was performed on the resulting bands visualized by ethidium-bromide agarose gel to quantify the intensity of the amplicons. The ratio of $A_0$ intensity to the intensity of H3.3 is plotted on the y-axis. The data used to generate this graph is shown in supplementary table S1.

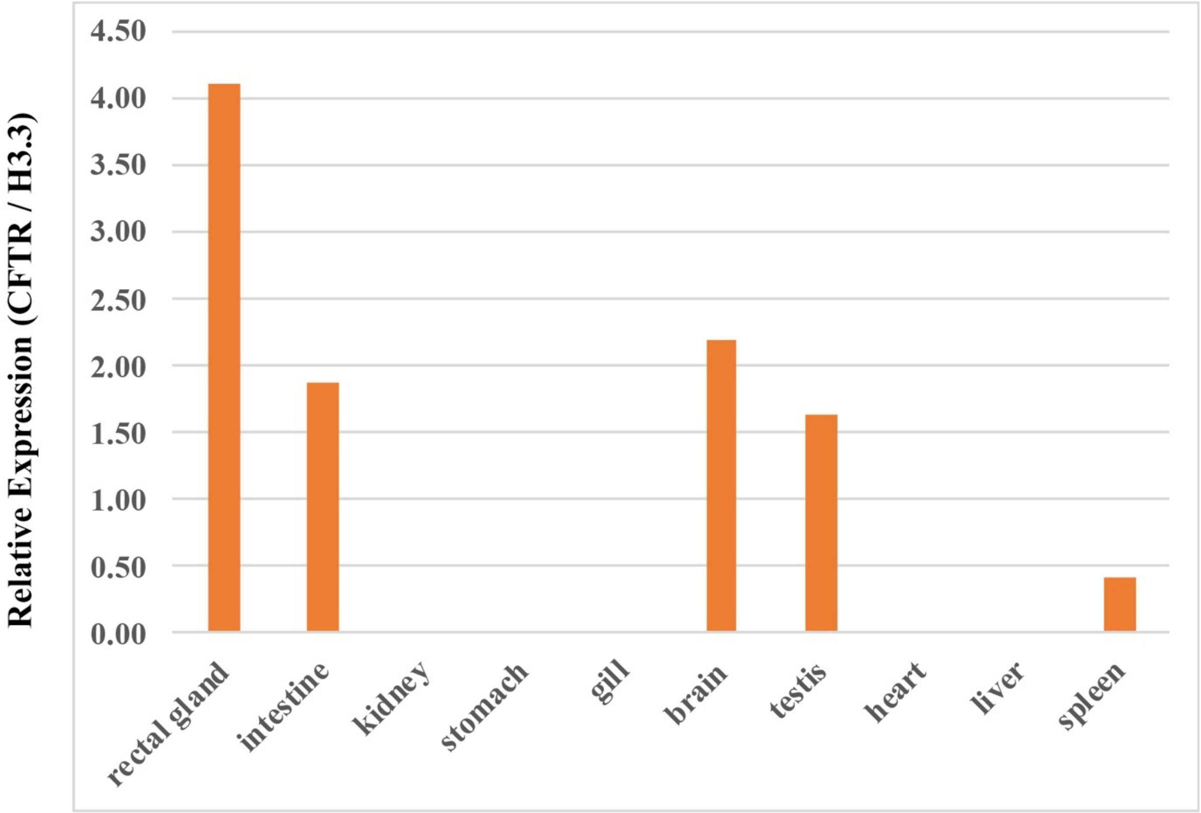

**Figure S4.** Expression of CFTR in selected shark tissues. Complimentary DNA (cDNA) was prepared from the indicated shark tissues and CFTR was amplified by quantitative RT-PCR using CFTR-specific primers. Shark histone H3.3 was also amplified simultaneously, and densitometry was performed on the resulting bands visualized by ethidium-bromide agarose gel to quantify the intensity of the amplicons. The ratio of CFTR intensity to the intensity of H3.3 is plotted on the y-axis. The data used to generate this graph is shown in supplementary table S2.

A

```
-322  aatcaccgcacagcaggcgacagcacttgggagggagggaggcaaaataaagaaattctgcaaggaagacatttaaaaaa
-242  aaataatgtcccacccagcaccagaggtcccgggacattctgtgcaaatgcttttgggaattcggtgaaagagaagtgat
-162  tgactctttctgctaaagctttgcgcatggcaaattgcaggaacgtcaagcagcaaagagaaaagaatgaaggagcaaag
 -82  aaggaactgagagataagaagaaggaaatgaagaaataagaagacaaagggagaggaagactacgcaattttttcatttca
  -2  cgATGGACATGTTTGAAAGTTTTGTCGATAACGTGGCAAACCTGTCGCACAACAAGATCAATTGCACCAGCACTTTCTTC
   1    M  D  M  F  E  S  F  V  D  N  V  A (N) L  S  H  N  K  I (N) C  T  S  T  F  F
                                                                I
  78  CACATCAAGATTGGGTACCTTATTATGGAGGTGTTAACGGCATTCTTTGCTGTGCTTGGGAACATCTTCATCTGTTTTGT
  27  H  I  K  I  G  Y  L  I  M  E  V  L  T  A  F  F  A  V  L  G  N  I  F  I  C  F  V
                                                                          II
 158  CGTCATCAGGAACAGGAAGCTGCGGACGGTGACCAATTACTTCCTGGTGTCGCTGGCCGTGGCGGACATACTGGTCGGGG
  54   V  I  R  N  R  K  L  R  T  V  T  N  Y  F  L  V  S  L  A  V  A  D  I  L  V  G  A

 238  CGGTGGCCATCCCGTGCGCCTTGCTGTCGAGCCTGGGCCTCCCCAGGTGCAGCTACTACCTCTGTGTCCTGATGCTGTGC
  81   V  A  I  P  C  A  L  L  S  S  L  G  L  P  R  C  S  Y  Y  L  C  V  L  M  L  C
                   III
 318  ACGCTGCTCGTGCTGACCCAAGCCTCCATCTTTGGGCTGTTCGCCATCGCTGTCGAGCGCTACATCGCCATCCTGACGCC
 107  T  L  L  V  L  T  Q  A  S  I  F  G  L  F  A  I  A  V  E  R  Y  I  A  I  L  T  P
                                                                           IV
 398  GTTCCGCTACCAGGCCTTGGTGACGTCCAGGAACGCGGGCCTGGTAATCGTCACTTCGTGGGTGCTGGCGGTCATCATCG
 134   F  R  Y  Q  A  L  V <T> S  R  N  A  G  L  V  I  V  T  S  W  V  L  A  V  I  I  G

 478  GGCTGGTGCCTCTGATGGGCTGGCGCAAAATTCCCATGGCCGACGAGAGATGCCTGTTCGATAACGTGATCGACGAGACC
 161   L  V  P  L  M  G  W  R  K  I  P  M  A  D  E  R  C  L  F  D  N  V  I  D  E  T
                                  V
 558  TACATGGTGTACTTCAATTTTATTTGTTGTATGCTCCTGCCTCTGCTCATCATGTTTGTCATCTACGGCAAGATATTTCT
 187  Y  M  V  Y  F  N  F  I  C  C  M  L  L  P  L  L  I  M  F  V  I  Y  G  K  I  F  L

 638  CGAGGTCAAGAAGCAGATCCGCCGAATAGCCGAGCGCCACATCAACATCAGCGCGGAGGAGAAGCGGCGGAAAATAATCC
 214   E  V  K  K  Q  I  R  R  I  A  E  R  H  I  N  I [S] A  E  E  K  R  R  K  I  I  R
                                                              VI
 718  GCAAGGAGGTCCAGACCGCCACGTCGCTCTTCATCGTGCTCTTCTGCTTCACCCTGAGCTGGATCCCCCTGCACATCCTG
 241   K  E  V  Q  T  A  T  S  L  F  I  V  L  F  C  F  T  L  S  W  I  P  L  H  I  L
                                                                             VII
 798  AACTGCGTCAAGCTCTCCTGTCCCAGCTGCGACATCCCGGCTTCCCTGATGCTGACCACGGTCATCCTGTCCCACATCAA
 267  N  C  V  K  L  S  C  P  S  C  D  I  P  A  S  L  M  L  T  T  V  I  L  S  H  I  N

 878  CTCGGTCGTCAACCCCATTGTCTACGTCTTCAGGATAAAGAGCTTCTGGGACGCTTTCGAAGACATCGTCTCCTGCCTCC
 294   S  V  V  N  P  I  V  Y  V  F  R  I  K [S] F  W  D  A  F  E  D  I  V  S  C  L  P

 958  CGTACCCCGCGGTCGTCGGAAAATATTCCAATGAGGATTTTACAGGGATGATTAAGATCAGGACATTCGGGAACAGGAAC
 321   Y  P  A  V  V  G  K  Y [S] N  E  D  F  T  G  M  I  K  I  R  T  F  G  N  R  N

1038  GATGCATTGCCAGGAACATAGatatgctgggagagatcctgtgtttacccagattggtgggtaaaccccacccttattaa
 347  D  A  L  P  G  T  *
1118  caattccaatttaccagacgcaatgacgcaccataactcgctctctatttttatgcttgctgtcaattgtacctctattt
1198  attgtacatttcc
```

B

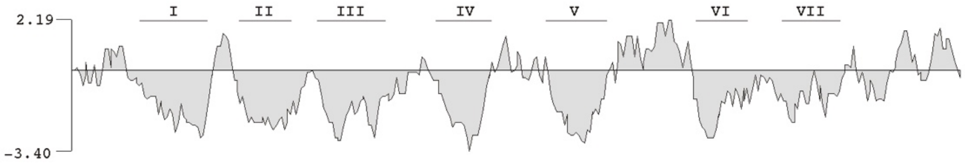

**Figure S5**. Nucleotide and deduced amino acid sequence of the shark $A_0$ adenosine receptor. (A) The seven predicted transmembrane segments (overlined), potential N-linked glycosylation sites (circles), potential casein kinase II phosphorylation sites (squares) and a potential protein kinase C phosphorylation site (diamond). (B) Kyte-Doolittle hydrophilicity plot (negative = more hydrophobic) with the predicted seven transmembrane domains indicated.

```
                                             *
Shark A0     1    MDMFESFVdnvanlshnkinctstffhikigYLIMEVLTAFFAVLGNIFICFVVIRNRKLRTVTNYFLVSLAVADILVGA   80
2ydv-struct  1    MPIMGSSV-----------------------YITVELAIAVLAILGNVLVCWAVWLNSNLQNVTNYFVVSAAAADILVGV   57

Shark A0     81   VAIPCALLSSLGLPRCSYYLCVLMLCTLLVLTQASIFGLFAIAVERYIAILTPFRYQALVTSRNAGLVIVTSWVLAVIIG  160
2ydv-struct  58   LAIPFAIAISTGFC-AACHGCLFIACFVLVLTASSIFSLLAIAIDRYIAIRIPLRYNGLVTGTRAKGIIAICWVLSFAIG  136

Shark A0     161  LVPLMGWRKIPMADE-------------RCLFDNVIDETYMVYFNFICCMLLPLLIMFVIYGKIFLEVKKQIRRIAERHI  227
2ydv-struct  137  LTPMLGWNNCGQPKEGKAHSQGCGEGQVACLFEDVVPMNYMVYFNFFACVLVPLLLMLGVYLRIFLAARRQLKQMESQPL  216

Shark A0     228  NISAEEKRRKIIRKEVQTATSLFIVLFCFTLSWIPLHILNCVKLSCPSCD-IPASLMLTTVILSHINSVVNPIVYVFRIK  306
2ydv-struct  217  ---PGERARSTLQKEVHAAKSLAIIVGLFALCWLPLHIINCFTFFCPDCSHAPLWLMYLAIVLSHTNSVVNPFIYAYRIR  293

Shark A0     307  SFWDAFEDIVSCLPYPAVVGKYSNEDFTGMIKIRTFGNrndalpgt  352
2ydv-struct  294  EFRQTFRKIIRSHVLR------QQEPFKAAAAENLYFQ--------  325
                                 *
```

**Figure S6.** Multiple amino acid sequence alignment between shark $A_0$ and human $A_{2a}$ receptor as extracted from the x-ray crystal structure of $A_{2a}$ deposited to the Protein Data Bank (PDB code 2ydv). The line labeled "2ydv-struct" represents the $A_{2a}$ sequence with disordered residues removed, but prior to the gap removal process depicted in supplementary fig. S7 below. The N-terminus of all adenosine receptors is the most variable in length and sequence, and therefore we chose to start the shark $A_0$ homology model at $A_0$ residue Tyr-32 (Y32) shown with an asterisk (*) above. For the C-terminus, we discovered that secondary structure prediction algorithms do not accurate predict the helical C-terminus of most receptors, as is particularly true in the case of 2ydv. We therefore chose a conservative truncation of the $A_0$ sequence for the homology model (also shown with asterisk above). NCBI Cobalt was used to construct the alignment and the red colored sequences represent the Cobalt 3-bit amino acid sequence similarity threshold.

**NCBI Cobalt Identity = 45% (red)**

```
A0 target    1     YLIMEVLTAFFAVLGNIFICFVVIRNRKLRTVTNYFLVSLAVADILVGAVAIPCALLSSLGLPCSYYLCVLMLCTLLVLT   80
A2a (2ydv)   1     YITVELAIAVLAILGNVLVCWAVWLNSNLQNVTNYFVVSAAAADILVGVLAIPFAIAISTGFCAACHGCLFIACFVLVLT   80

A0 target    81    QASIFGLFAIAVERYIAILTPFRYQALVTSRNAGLVIVTSWVLAVIIGLVPLMGWRKIPMADERCLFDNVIDETYMVYFN   160
A2a (2ydv)   81    ASSIFSLLAIAIDRYIAIRIPLRYNGLVTGTRAKGIIAICWVLSFAIGLTPMLGWNNCGQPKEACLFEDVVPMNYMVYFN   160

A0 target    161   FICCMLLPLLIMFVIYGKIFLEVKKQIRRIAERHIIRKEVQTATSLFIVLFCFTLSWIPLHILNCVKLSCPSCDIPASLM   240
A2a (2ydv)   161   FFACVLVPLLLMLGVYLRIFLAARRQLKQMESQSTLQKEVHAAKSLAIIVGLFALCWLPLHIINCFTFFCPDCSAPLWLM   240

A0 target    241   LTTVILSHINSVVNPIVYVFRIKSFWDAFEDIVSCLPY   278
A2a (2ydv)   241   YLAIVLSHTNSVVNPFIYAYRIREFRQTFRKIIRSHVL   278
```

**NCBI Cobalt 3-bit similarity = 81% (red)**

```
A0 target    1     YLIMEVLTAFFAVLGNIFICFVVIRNRKLRTVTNYFLVSLAVADILVGAVAIPCALLSSLGLPCSYYLCVLMLCTLLVLT   80
A2a (2ydv)   1     YITVELAIAVLAILGNVLVCWAVWLNSNLQNVTNYFVVSAAAADILVGVLAIPFAIAISTGFCAACHGCLFIACFVLVLT   80

A0 target    81    QASIFGLFAIAVERYIAILTPFRYQALVTSRNAGLVIVTSWVLAVIIGLVPLMGWRKIPMADERCLFDNVIDETYMVYFN   160
A2a (2ydv)   81    ASSIFSLLAIAIDRYIAIRIPLRYNGLVTGTRAKGIIAICWVLSFAIGLTPMLGWNNCGQPKEACLFEDVVPMNYMVYFN   160

A0 target    161   FICCMLLPLLIMFVIYGKIFLEVKKQIRRIAERHIIRKEVQTATSLFIVLFCFTLSWIPLHILNCVKLSCPSCDIPASLM   240
A2a (2ydv)   161   FFACVLVPLLLMLGVYLRIFLAARRQLKQMESQSTLQKEVHAAKSLAIIVGLFALCWLPLHIINCFTFFCPDCSAPLWLM   240

A0 target    241   LTTVILSHINSVVNPIVYVFRIKSFWDAFEDIVSCLPY   278
A2a (2ydv)   241   YLAIVLSHTNSVVNPFIYAYRIREFRQTFRKIIRSHVL   278
```

**Figure S7.** NCBI Cobalt alignment of shark $A_0$ target sequence and $A_{2a}$ template sequence with "gap" sequences present in one receptor but not the other removed. The top alignment shows simply amino acid identity in red, and the bottom alignment shows Cobalt 3-bit amino acid sequence similarly threshold, also in red. The alignment was used as a guide to manually mutate the $A_{2a}$ sequence into $A_0$ using PyMOL taking into consideration rotameric library preferences for each side chain and the energy minimization of clashes with the existing structure.

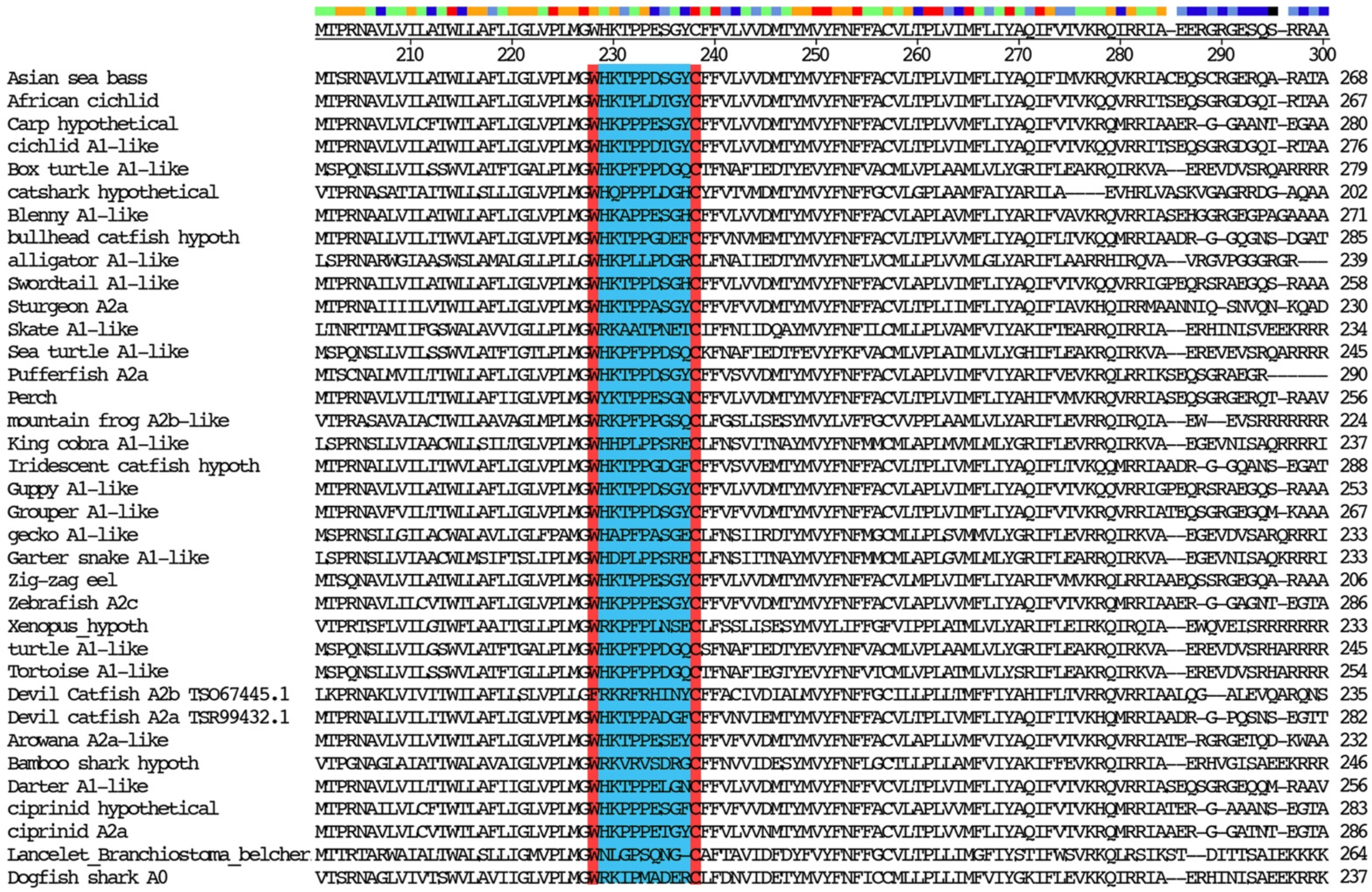

**Figure S8**. Adenosine receptor ancestors exhibit a minimally short extracellular loop 2 (ECL2) which is exactly 9 amino acid residues (highlighted in blue) between invariant tryptophan (W) and cysteine (C) markers, which are highlighted in red. Ancestor adenosine receptors appear limited to fish, amphibians and reptiles, as we could find none in birds or mammals. The primitive lancelet has an exceptionally short ECL2, being 1 residue shorter than the rest. This putative receptor remains uncharacterized, but since it contains all of the adenosine-interacting motifs we have included it in this alignment. Note that the lancelet putative ADOR is also ancestral with respect to all $A_{2a}$- and $A_{2b}$- ADORs (see figure 1 of the main text).

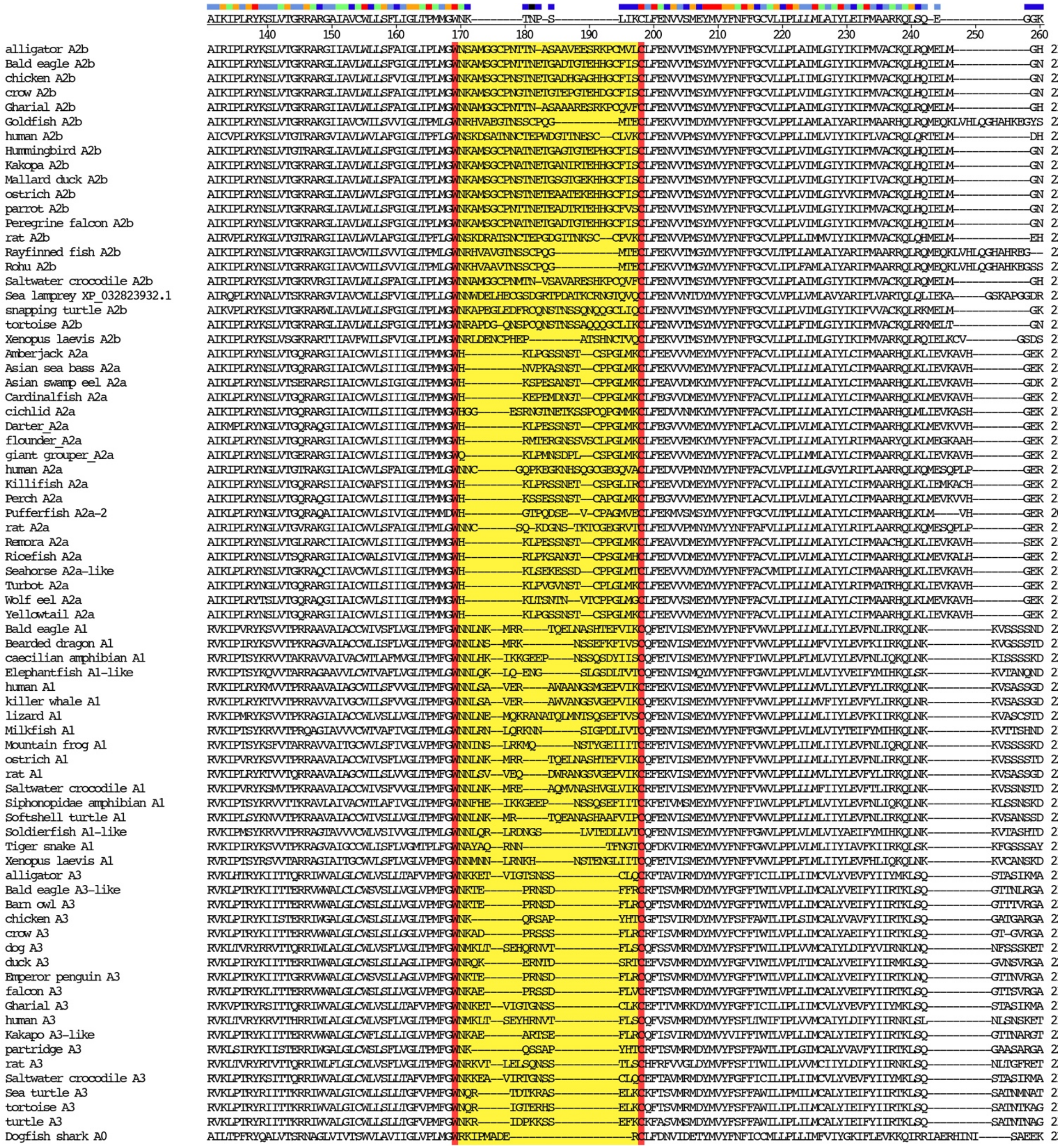

**Figure S9**. Specialized ADORs exhibit an elongated ECL2 with respect to ancestor receptors. The tryptophan (W) and cysteine (C) markers used in Figure S8 are used here (in red) to measure $ECL2_{length}$ (yellow). For reference, the shark $A_0$ receptor ancestor ($ECL2_{length}$=9) is shown at the bottom. Selected specialized ADORs from mammals, birds, reptiles and fish are shown and the total receptors analyzed in the main alignment not shown were: $A_1$ (n=46), $A_{2a}$ (n=23), $A_{2b}$ (n=30) and $A_3$ (n=18). Every specialized receptor analyzed (n=117 total) had $ECL2_{length} \geq 10$ aa. Avian $A_3$ receptors from partridge and chicken have a 10-aa ECL2, and the next longest ECL2 (at 12-aa) were found in other avian $A_3$. $A_{2b}$ receptors contain the longest ECL2 (from 19-41aa). Not only were we surprised to find specialized ADORs in the whale shark ($A_1$ and $A_{2b}$), the carpet shark ($A_1$) and the skate ($A_1$, $A_{2a}$ and $A_{2b}$) but these ADORs had the longest ECL2 of any of their counterparts in other species (not shown).

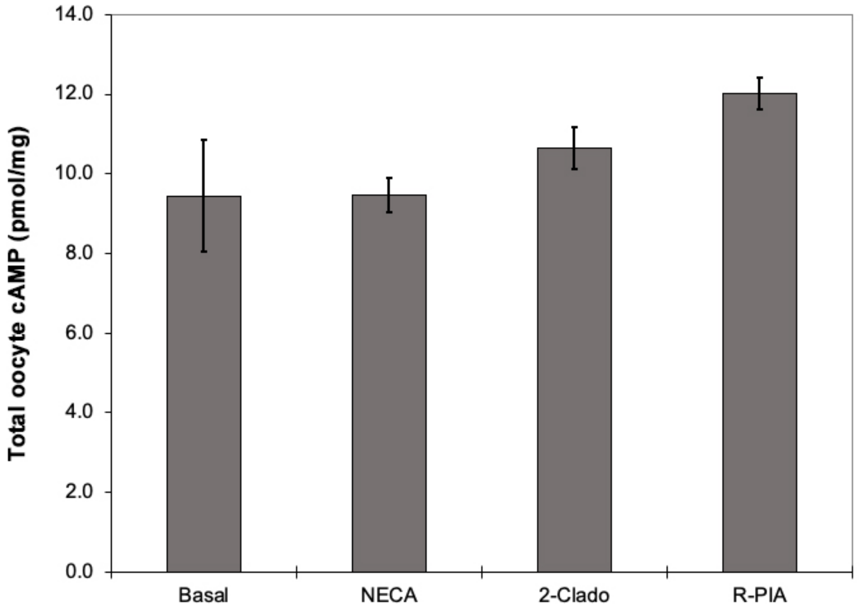

**Figure S10**. Total cAMP in whole oocytes. Oocytes injected with the shark $A_0$ receptor only were subjected to agonists (NECA, 2-ClAdo or R-PIA, 10 µM each) or Frog Ringer's ("Basal"). Cyclic AMP was measured by radioimmunoassay as described in Supplemental References (1). Bars are plotted as the means ± SD (n=5 oocytes for each condition). Probabilities (basal vs agonist, agonist1 vs agonist2, etc.) were calculated using the Student's two-tailed homoscedastic t-Test. Due to the high SD of the Basal condition, only when R-PIA was compared to NECA ($p=0.001$) or 2-ClAdo ($p=0.047$) were values deemed statistically significant ($p<0.05$).

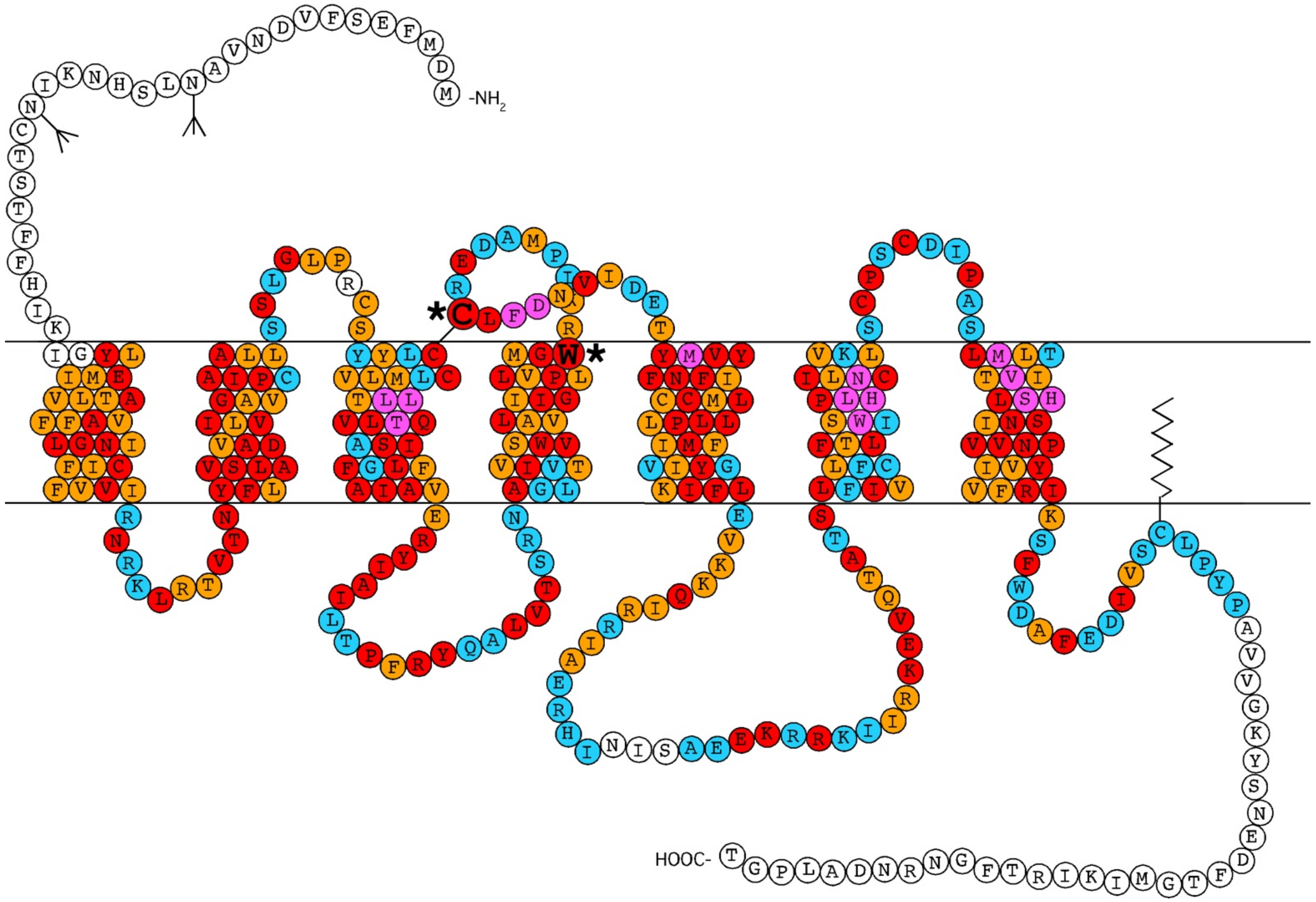

**Figure S11**. Topological model of the shark $A_0$ adenosine receptor and similarity to human ADORs. Potential glycosylation sites in the N-terminal section are indicated. Amino acids colored red are identical to the human $A_1$- or $A_{2a}$- adenosine receptors, those colored orange have high chemical similarity, and those colored blue are distinct. Amino acids in magenta are either identical to- or highly similar to- known adenosine-interacting residues based on the high-resolution x-ray crystal structure of the human $A_{2a}$ receptor (PDB code 2ydo) and the cryo-EM structure of the human $A_1$ receptor (PDB code 6d9h). All colored amino acid residues have good backbone correspondence of human ADORs to our shark $A_0$ homology model core. Two invariantly conserved amino acids (aa) in 166 receptors analyzed (marked with asterisk) served as a marker for establishing the evolution of extracellular loop 2 (ECL2). Ancestral adenosine receptors exhibited a minimally short ECL2 of 9 amino acids, and all of the specialized adenosine receptor subtypes examined have an elongated ECL2 ($A_1$ elongated by 4-17 aa, $A_{2a}$ elongated by 8-14 aa, $A_{2b}$ elongated by 10-19 aa and $A_3$ elongated by 1-7 aa). A minimally short ECL2 emerges as a structural characteristic of all ancestral adenosine receptors (see multiple sequence alignments of ECL2 in supplementary figs. S8 and S9).

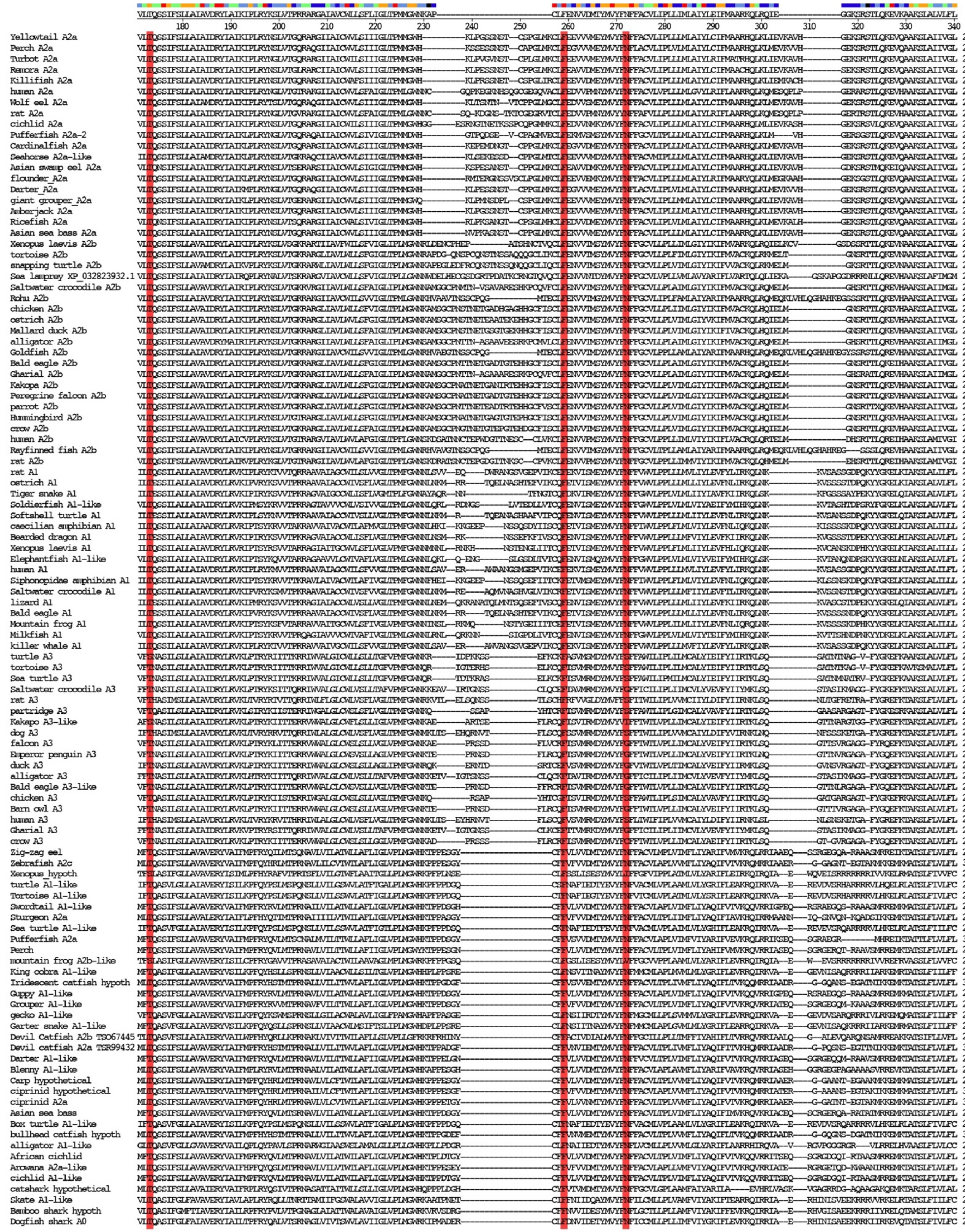

**Figure S12**. Page 1 of multi-alignment of 110 selected specialized- or ancestral- receptor subtypes. Conserved amino acid residues involved in adenosine- or adenosine analog- binding are highlighted in red. $A_0$ T112, **F179** (invariant aromatic interaction with purine ring) and **N192** (invariant two hydrogen bonds with purine ring) are highlighted in red.

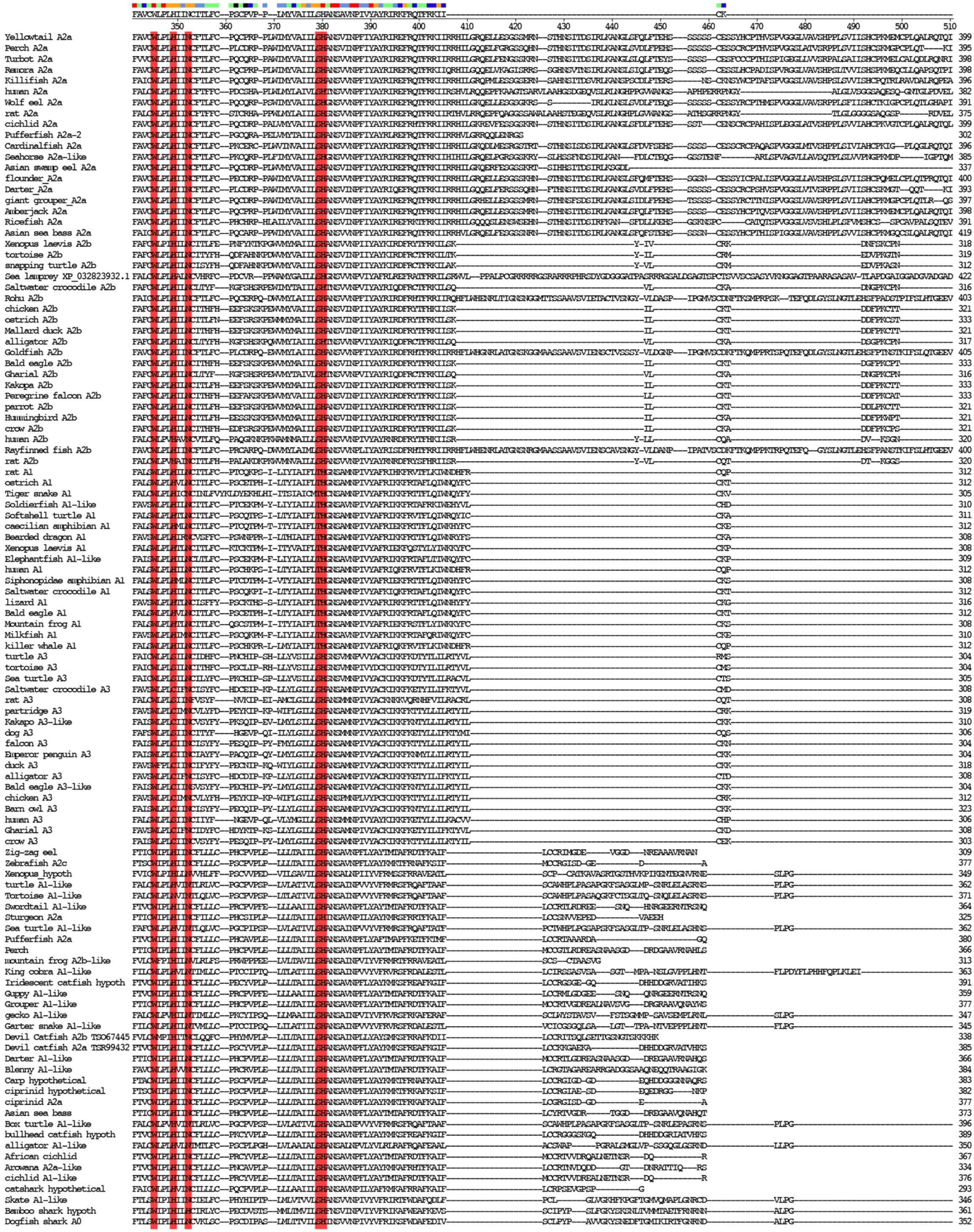

**Figure S13**. Page 2 of multi-alignment of 110 selected specialized- or ancestral- receptor subtypes. Conserved amino acid residues involved in adenosine- or adenosine analog- binding are highlighted in red. $A_0$ W260, H264, N267 and S290/H291 are highlighted in red.

**Table S1.** Results of densitometry scanning of PCR amplicons for shark $A_0$ receptor and histone H3.3 from cDNA obtained from total RNA prepared from different shark tissues. The ratio of shark $A_0$ receptor expression to H3.3 expression is shown in the rightmost column for each tissue, and this column is plotted in the graph shown in fig. S3.

| **Source of cDNA** | **integrated intensity $A_0$** | **integrated intensity H3.3** | **relative amount $A_0$ receptor expression** |
|---|---|---|---|
| rectal gland | 598.2 | 203.4 | 2.94 |
| intestine | 8.3 | 68.9 | 0.12 |
| kidney | 239.8 | 308.7 | 0.78 |
| stomach | 511.5 | 133.0 | 3.85 |
| gill | 9.0 | 255.4 | 0.04 |
| brain | 20.8 | 51.7 | 0.40 |
| testis | 48.7 | 268.2 | 0.18 |
| heart | 29.8 | 228.4 | 0.13 |
| liver | 37.8 | 267.7 | 0.14 |
| spleen | 20.7 | 50.5 | 0.41 |

**Table S2.** Results of densitometry scanning of PCR amplicons for shark CFTR and histone H3.3 from cDNA obtained from total RNA prepared from different shark tissues. The ratio of shark CFTR expression to H3.3 expression is shown in the rightmost column for each tissue, and this column is plotted in the graph shown in fig. S4.

| **Source of cDNA** | **integrated intensity CFTR** | **integrated intensity H3.3** | **relative amount CFTR expression** |
|---|---|---|---|
| rectal gland | 438.6 | 106.8 | 4.11 |
| intestine | 62.1 | 33.2 | 1.87 |
| kidney | 0.0 | 112.1 | 0.00 |
| stomach | 0.0 | 16.2 | 0.00 |
| gill | 0.0 | 39.2 | 0.00 |
| brain | 341.9 | 155.9 | 2.19 |
| testis | 227.2 | 138.2 | 1.64 |
| heart | 0.0 | 75.7 | 0.00 |
| liver | 0.0 | 152.4 | 0.00 |
| spleen | 6.0 | 14.3 | 0.42 |

**Table S3 (multiple tables on the following pages)**

| | | | | |
|---|---|---|---|---|
| | **Experiment#** | 1 | **Date** | February 27 |
| | **cRNA:** | 10ng A0-ADOR & 5% CFTR | | |
| | **Days after inj:** | 4 | **For 5μM:** | 4850 nA |
| **No** | **Agonist** | **Rel** | **Agonist** | **Effect on I** |
| 1 | S-PIA | > | CGS21680 | Decrease |
| 2 | S-PIA | > | CGS21680 | Increase |
| 3 | S-PIA | > | NECA | Decrease |
| 4 | CGS21680 | > | NECA | Increase |
| 5 | NECA | >?? | CGS21680 | Increase |
| 6 | S-PIA | > | NECA | Increase |
| 7 | S-PIA | > | NECA | Decrease |
| 8 | S-PIA | > | NECA | Decrease |
| 9 | CPA | > | NECA | Increase |
| **Result: S-PIA > CGS21680 > NECA** | | | | |
| **Result: CPA > NECA** | | | | |

**Summaries of experiments performed in 10 different oocytes (see also following pages). Potency response to compounds were measured in pairs at equivalent concentrations, and the relative potency determined by either an increase- or decrease in total chloride current from the oocyte in response to the second compound. Oocyte response to the first compound was always allowed to stabilize before perifusion of the second compound (which lacked the first compound). Pairs of compounds were often applied to the oocyte in both orders to determine reproducibility within each experiment. For example, in Experiment #1, steps 1 and 2, the agonists S-PIA and CGS21680 were applied to the oocyte in opposite orders, which had an opposite effect on the total current from the oocyte. For clarity, all tables were prepared placing the most potent compound of the pair (agonist or antagonist) in the leftmost column. The overall potency order for all compounds tested in each experiment are summarized at the bottom of each table. At the end of each experiment, the health of the oocyte was tested with the application of 5 μM forskolin, and the total current was recorded in the header of each table.**

| | | | | |
|---|---|---|---|---|
| | **Experiment#** | 2 | **Date** | February 28 |
| | **cRNA:** | 15ng A0-ADOR & 5% CFTR | | |
| | **Days after inj:** | 5 | **For 5μM:** | 4600 nA |
| **No** | **Agonist** | **Rel** | **Agonist** | **Effect on I** |
| 1 | S-PIA | > | CGS21680 | Increase |
| 2 | S-PIA | > | NECA | Decrease |
| 3 | CGS21680 | > | NECA | Increase |
| 4 | CGS21680 | > | NECA | Decrease |
| 5 | CGS21680 | > | NECA | Increase |
| 6 | S-PIA | > | CGS21680 | Increase |
| 7 | S-PIA | > | 2ClAdo | Decrease |
| 8 | CGS21680 | > | 2ClAdo | Increase |
| 9 | S-PIA | > | CGS21680 | Increase |
| 10 | S-PIA | > | 2ClAdo | Decrease |
| 11 | CGS21680 | > | 2ClAdo | Increase |
| 12 | CGS21680 | > | 2ClAdo | Decrease |
| 13 | 2ClAdo | > | NECA | Decrease |
| 14 | CGS21680 | > | 2ClAdo | Increase |
| 15 | S-PIA | > | CGS21680 | Increase |
| 16 | R-PIA | > | S-PIA | Increase |
| 17 | R-PIA | > | CPA | Decrease |
| 18 | R-PIA | > | CPA | Increase |
| 19 | R-PIA | > | S-PIA | Decrease |
| **Result: R-PIA > S-PIA > CGS21680 > 2ClAdo > NECA** | | | | |
| **Result: R-PIA > CPA** | | | | |

| | | | | | |
|---|---|---|---|---|---|
| | | **Experiment#** | 3 | **Date** | March 26 |
| | | **cRNA:** | 15ng A0-ADOR & 2.5% CFTR | | |
| | | **Days after inj:** | 4 | **For 5$\mu$M:** | 535 nA |
| | **No** | **Agonist** | **Rel** | **Agonist** | **Effect on I** |
| | 1 | 2ClAdo | > | NECA | Increase |
| | 2 | CGS21680 | > | 2ClAdo | Increase |
| | 3 | CGS21680 | > | CV1808 | Decrease |
| | 4 | CV1808 | > | NECA | Decrease |
| | 5 | 2ClAdo | > | NECA | Increase |
| | 6 | 2ClAdo | > | CV1808 | Decrease |
| | 7 | CV1808 | > | DPMA | Decrease |
| | 8 | DPMA | > | NECA | Decrease |
| | 9 | R-PIA | > | CPA | Decrease |
| | 10 | S-PIA | > | CPA | Increase |
| | 11 | R-PIA | > | S-PIA | Increase |
| | 12 | R-PIA | > | S-PIA | Decrease |
| | 13 | S-PIA | > | CPA | Decrease |
| | 14 | CGS21680 | > | CPA | Increase |
| | 15 | S-PIA | > | CGS21680 | Increase |
| | 16 | S-PIA | > | CGS21680 | Decrease |
| | 17 | CGS21680 | > | CPA | Decrease |
| | 18 | CPA | > | 2ClAdo | Decrease |
| | 19 | CV1808 | >? | 2ClAdo | Increase |
| | 20 | CV1808 | > | 2ClAdo | Decrease |
| | 21 | CPA | > | 2ClAdo | Increase |
| | 22 | CPA | > | CV1808 | Decrease |
| | 23 | CV1808 | > | 2ClAdo | Decrease |
| | 24 | DPMA | > | 2ClAdo | Increase |
| | 25 | CV1808 | > | DPMA | Increase |
| | 26 | CV1808 | > | DPMA | Decrease |
| | 27 | DPMA | > | 2ClAdo | Decrease |
| | 28 | NECA | > | 2ClAdo | Decrease |
| | 29 | NECA | > | 2ClAdo | Increase |
| | 30 | NECA | > | 2ClAdo | Decrease |
| **Result: R-PIA > S-PIA > CGS21680 > CPA > 2ClAdo > CV1808 > DPMA > NECA** | | | | | |

| | | | | |
|---|---|---|---|---|
| | **Experiment#** | 4 | **Date** | March 26 |
| | **cRNA:** | 15ng A0-ADOR & 2.5% CFTR | | |
| | **Days after inj:** | 4 | **For 5μM:** | 345 nA |
| **No** | **Agonist** | **Rel** | **Agonist** | **Effect on I** |
| 1 | 2ClAdo | > | NECA | Decrease |
| 2 | 2ClAdo | > | NECA | Increase |
| 3 | 2ClAdo | > | CV1808 | Decrease |
| 4 | CV1808 | > | NECA | Decrease |
| 5 | CV1808 | > | NECA | Increase |
| 6 | 2ClAdo | > | CV1808 | Increase |
| 7 | 2ClAdo | > | DPMA | Decrease |
| 8 | CV1808 | > | DPMA | Decrease |
| 9 | 2ClAdo | > | CV1808 | Increase |
| 10 | CPA | > | 2ClAdo | Increase |
| 11 | CGS21680 | >?? | CPA | Increase |
| 12 | S-PIA | > | CGS21680 | Increase |
| 13 | R-PIA | > | S-PIA | Increase |
| 14 | R-PIA | > | S-PIA | Decrease |
| 15 | S-PIA | > | CGS21680 | Decrease |
| 16 | CPA | > | DPMA | Decrease |
| 17 | 2ClAdo | > | CV1808 | Decrease |
| 18 | CV1808 | > | NECA | Decrease |
| 19 | 8PT | > | CGS15943 | Increase |
| 20 | 8PT | > | CGS15943 | Decrease |
| 21 | 8PT | ?? | CGS15943 | |
| 22 | 8PT | > | CSC | Decrease |
| 23 | 8PT | > | CSC | Increase |
| **Result: R-PIA > S-PIA > CGS21680 > CPA > 2ClAdo > CV1808 > DPMA > NECA** | | | | |
| **Result (Antagonist): 8PT > CGS15943** | | | | |
| **Result (Antagonist): 8PT > CSC** | | | | |

| | | | | |
|---|---|---|---|---|
| | **Experiment#** | 5 | **Date** | March 27 |
| | **cRNA:** | 15ng A0-ADOR & 2.5% CFTR | | |
| | **Days after inj:** | 4 | **For 5μM:** | 150 nA |
| **No** | **Agonist** | **Rel** | **Agonist** | **Effect on I** |
| 1 | 2ClAdo | > | NECA | Decrease |
| 2 | R-PIA | > | NECA | Increase |
| 3 | R-PIA | > | S-PIA | Decrease |
| 4 | S-PIA | > | CGS21680 | Decrease |
| 5 | CPA | > | CGS21680 | Increase |
| 6 | CPA | > | CV1808 | Decrease |
| | DPMA | ???? | CV1808 | |
| 7 | CV1808 | >?? | DPMA | Increase |
| 8 | DPMA | >?? | CV1808 | Increase |
| 9 | R-PIA | > | S-PIA | |
| 10 | CGS15943 | > | 8PT | Increase |
| 11 | PD115199 | > | CGS15943 | Increase |
| 12 | PD115199 | > | CGS15943 | Decrease |
| 13 | DPCPX | > | CGS15943 | Decrease |
| 14 | DPCPX | >?? | PD115199 | |
| 15 | DPCPX | > | CSC | Decrease |
| 16 | CSC | > | CGS15943 | Decrease |
| 17 | 8PT | >?? | CGS15943 | Increase |
| 18 | CSC | > | 8PT | Decrease |
| **Result: R-PIA > S-PIA > CPA > CGS21680 > CV1808 > DPMA** | | | | |
| **Result: 2ClAdo > NECA** | | | | |
| **Result (Antagonist): DPCPX > PD115199 > CGS15943 > 8PT** | | | | |
| **Result (Antagonist): CSC > CGS15943** | | | | |

| | | | | |
|---|---|---|---|---|
| | **Experiment#** | 6 | **Date** | April 4 |
| | **cRNA:** | 15ng A0-ADOR & 2.5% CFTR | | |
| | **Days after inj:** | 5 | **For 5μM:** | 750 nA |
| **No** | **Agonist** | **Rel** | **Agonist** | **Effect on I** |
| 1 | DPMA | > | NECA | Increase |
| 2 | DPMA | >?? | CV1808 | Decrease |
| 3 | CV1808 | >?? | DPMA | Decrease |
| 4 | DPMA | > | NECA | Decrease |
| 5 | DPMA | > | NECA | Increase |
| 6 | DPMA | >?? | CV1808 | Decrease |
| 7 | 2ClAdo | > | CV1808 | Increase |
| 8 | CPA | > | 2ClAdo | Increase |
| 9 | CPA | > | CGS21680 | Decrease |
| 10 | CPA | > | CGS21680 | Increase |
| 11 | CPA | > | CGS21680 | Decrease |
| 12 | S-PIA | > | CGS21680 | Increase |
| 13 | R-PIA | > | S-PIA | Increase |
| 14 | 8PT | > | CSC | Increase |
| 15 | 8PT | >? | CGS15943 | Decrease |
| **Result: R-PIA > S-PIA** | | | | |
| **Result: CPA > CGS21680 > 2ClAdo > CV1808 > DPMA > NECA** | | | | |
| **Result (Antagonist): 8PT > CSC** | | | | |
| **Result (Antagonist): 8PT > CGS15943** | | | | |

| | Experiment# | 7 | Date | April 5 |
|---|---|---|---|---|
| | cRNA: | 15ng A0-ADOR & 2.5% CFTR | | |
| | Days after inj: | 4 | For 5μM: | 2300 nA |
| No | Agonist | Rel | Agonist | Effect on I |
| 1 | CV1808 | > | NECA | Increase |
| 2 | 2ClAdo | > | CV1808 | Increase |
| 3 | CV1808 | > | DPMA | Decrease |
| 4 | DPMA | > | NECA | Decrease |
| 5 | CV1808 | > | NECA | Increase |
| 6 | CV1808 | > | DPMA | Decrease |
| 7 | DPMA | > | NECA | Increase |
| 8 | CV1808 | > | DPMA | Increase |
| 9 | 2ClAdo | > | CV1808 | Increase |
| 10 | CPA | > | 2ClAdo | Increase |
| Result: CPA > 2ClAdo > CV1808 > DPMA > NECA | | | | |

| | Experiment# | 8 | Date | April 19 |
|---|---|---|---|---|
| | cRNA: | 15ng A0-ADOR & 2.5% CFTR | | |
| | Days after inj: | 4 | For 5μM: | not recorded |
| No | Agonist | Rel | Agonist | Effect on I |
| 1 | DPMA | > | NECA | Increase |
| 2 | CV1808 | > | DPMA | Increase |
| 3 | DPMA | > | CV1808 | Decrease |
| 4 | DPMA | >? | CV1808 | Increase |
| 5 | DPMA | > | CV1808 | Decrease |
| 6 | DPMA | >? | CV1808 | Increase |
| 7 | 2ClAdo | > | DPMA | Increase |
| 8 | CPA | > | 2ClAdo | Increase |
| 9 | CPA | > | CGS21680 | Decrease |
| 10 | CPA | > | CGS21680 | Decrease |
| 11 | S-PIA | > | CPA | Increase |
| 12 | R-PIA | > | S-PIA | Increase |
| 13 | CV1808 | > | NECA | Increase |
| 14 | DPMA | > | CV1808 | Increase |
| 15 | 2ClAdo | > | DPMA | Increase |
| 16 | CGS21680 | > | 2ClAdo | Increase |
| 17 | CPA | > | CGS21680 | Increase |
| 18 | S-PIA | >?? | CPA | Increase |
| 19 | CPA | > | S-PIA | Decrease |
| 20 | R-PIA | > | S-PIA | Increase |
| 21 | CGS15943 | >?? | 8PT | Decrease |
| 22 | 8PT | > | CGS15943 | Decrease |
| 23 | CSC | > | CGS15943 | Increase |
| 24 | 8PT | > | CSC | Increase |
| 25 | PD115199 | > | 8PT | Increase |
| 26 | DPCPX | > | PD115199 | Increase |
| 27 | DPCPX | > | CGS15943 | Decrease |
| 28 | CSC | > | CGS15943 | Increase |
| 29 | 8PT | > | CSC | Increase |
| 30 | PD115199 | > | 8PT | Increase |
| Result: CPA > R-PIA > S-PIA > CGS21680 > 2ClAdo > DPMA > CV1808 > NECA | | | | |
| Result: DPMA > CV1808 | | | | |
| Result: CPA > S-PIA | | | | |
| Result (Antagonist): DPCPX > PD115199 > 8PT > CSC > CGS15943 | | | | |

| | | | | |
|---|---|---|---|---|
| | **Experiment#** | 9 | **Date** | April 23 |
| | **cRNA:** | 15ng A0-ADOR & 2.5% CFTR | | |
| | **Days after inj:** | 4 | **For 5μM:** | not recorded |
| **No** | **Agonist** | **Rel** | **Agonist** | **Effect on I** |
| 1 | CGS15943 | > | CSC | Decrease |
| 2 | 8PT | > | CSC | Increase |
| 3 | PD115199 | > | 8PT | Increase |
| 4 | DPCPX | > | PD115199 | Decrease |
| 5 | DPCPX | > | PD115199 | Decrease |
| 6 | 8PT | > | CGS15943 | Increase |
| 7 | 8PT | > | CGS15943 | Decrease |
| 8 | CSC | > | CGS15943 | Decrease |
| 9 | CSC | > | CGS15943 | Increase |
| **Result (Antagonist): DPCPX > PD115199 > 8PT > CSC > CGS15943** | | | | |
| | **Result (Antagonist): CGS15943 > CSC** | | | |

| | | | | |
|---|---|---|---|---|
| | **Experiment#** | 10 | **Date** | May 4 |
| | **cRNA:** | 15ng A0-ADOR & 2.5% CFTR | | |
| | **Days after inj:** | 5 | **For 5μM:** | 1733 nA |
| **No** | **Agonist** | **Rel** | **Agonist** | **Effect on I** |
| 1 | DPMA | > | NECA | Increase |
| 2 | 2ClAdo | > | DPMA | Increase |
| 3 | 2ClAdo | >? | CGS21680 | Decrease |
| 4 | 2ClAdo | >? | CGS21680 | Increase |
| 5 | 2ClAdo | > | CGS21680 | Decrease |
| 6 | CGS21680 | > | DPMA | Decrease |
| 7 | DPMA | > | NECA | Decrease |
| 8 | DPMA | > | NECA | Increase |
| 9 | DPMA | > | CV1808 | Decrease |
| 10 | DPMA | > | CV1808 | Increase |
| 11 | 2ClAdo | > | DPMA | Increase |
| 12 | CPA | ?? | 2ClAdo | |
| 13 | CPA | > | CGS21680 | Decrease |
| 14 | 2ClAdo | > | CGS21680 | Decrease |
| 15 | CPA | > | 2ClAdo | Increase |
| 16 | CPA | > | S-PIA | Decrease |
| 17 | R-PIA | > | S-PIA | Increase |
| 18 | R-PIA | > | CPA | Decrease |
| 19 | CPA | > | S-PIA | Decrease |
| 20 | **2ClAdo** | > | **S-PIA** | Increase |
| 21 | new-S-PIA | > | 2ClAdo | Increase |
| 22 | CPA | > | new-S-PIA | Increase |
| 23 | CPA | > | new-S-PIA | Decrease |
| **Result: R-PIA > CPA > S-PIA > 2ClAdo > CGS21680 > DPMA > CV1808 > NECA** | | | | |

**Table S4.** Comparison of the pharmacological profiles determined in the *Xenopus* oocyte (this work) to available functional and binding studies on Human Adenosine Receptors.

**Human $A_1$ agonist Profile**

| | |
|---|---|
| Human $A_1$ oocyte | CPA>R-PIA=NECA>2ClAdo>S-PIA>DPMA>CV1808>CGS21680 |
| Human $A_1$ Binding Study | R-PIA>NECA>Ado>S-PIA>>CGS21680 |
| Human $A_1$ Functional studies | CPA>R-PIA=CHA≥NECA>2ClAdo>S-PIA>CV1808≥CGS21680 |

**Human $A_{2b}$ agonist Profile**

| | |
|---|---|
| Human $A_{2b}$ oocyte | NECA>R-PIA>2ClAdo>CPA>DPMA>S-PIA>CV1808>CGS21680 |
| Human $A_{2b}$ Binding Study | Profile unavailable |
| Human $A_{2b}$ Functional Studies | NECA>2ClAdo>R-PIA=CHA>S-PIA≥CV1808≥CGS21680 |

**Human $A_{2b}$ antagonist Profile**

| | |
|---|---|
| Human $A_{2b}$ oocyte | 8PT>PD115199>DPCPX>CGS15943>CSC |
| Human $A_{2b}$ Binding Study | Inhibited by theophylline |
| Human A2b Functional Studies | DPCPX=8PT≥PD115199 |
| Rat Binding Studies | CGS15943>PD115199>CP66713>XAC>CPX>8PT>Alloxazine>8-p-SPT>HTQZ>Tracazolate>CSC>DPMX>Caffeine>Theophylline |

**Table S5.** Geometric properties of the refined shark $A_0$ homology model compared to structures of human ADORs.

| | **A0** | **A2a (2ydv)** | **A1 (6d9h)** |
|---|---|---|---|
| Ramachandran favored (%) | 92.74 | 97.43 | 96.58 |
| Ramachandran allowed (%) | 7.26 | 2.57 | 3.42 |
| Ramachandran outliers (%) | 0.00 | 0.00 | 0.00 |
| Rotamer outliers (%) | 0.36 | 1.92 | 0.00 |
| C-beta deviations | 0 | 0 | 0 |
| MolProbity score | 2.24 | 1.61 | 1.42 |

**Table S6**. Nine highly conserved- or invariant- amino acid residues are present in shark $A_0$ and are in close proximity to potentially interact with ligand in our $A_0$ homology model.

| A0 | A2a | A1 | reference |
|---|---|---|---|
| T112 | T88 | T91 | (Jiang, et al. 1996) |
| F179 | F168 | F171 | (Lane, et al. 2012) |
| D180 | E169 | E172 | (Kim, et al. 1996) |
| N192 | N181 | N184 | (Lebon, et al. 2011) |
| W260 | W246 | W247 | (Lane, et al. 2012) |
| H264 | H250 | H251 | (Jiang, et al. 1997) |
| N267 | N253 | N254 | (Lane, et al. 2012) |
| S290 | S277 | T277 | (Lane, et al. 2012) |
| H291 | H278 | H278 | (Askalan and Richardson 1994) |

**Table S7.** A1 adenosine receptor agonist and antagonist profiles in the literature.

| Subtype | Ago/Ant | Species | Tissue | Profile | Reference |
|---|---|---|---|---|---|
| A1 | Agonist | Review of functional studies | | CPA>R-PIA=CHA≥NECA>2ClAdo>S-PIA>CV1808≥CGS21680 | (2) |
| A1 | Agonist | Human | Brain | R PIA>NECA>Ado>S PIA»CGS21680 | (3) |
| A1 | Agonist | Human | Brain | CPA≥CHA>R-PIA≥2ClAdo≥NECA>S-PIA | (4) |
| A1 | Agonist | Bovine | Brain | R-PIA>S-PIA>NECA | (5) |
| A1 | Agonist | Bovine | Brain | R-PIA>CHA>CPA>S-PIA»NECA>2ClAdo | (6) |
| A1 | Agonist | Guinea pig | Brain | CPA»CCPA>R-PIA>CHA»NECA>Ado>BA>CGS21680 | (7) |
| A1 | Agonist | Rat | Testis | CPA>R-PIA>NECA>S-PIA | (8) |
| A1 | Agonist | Rat | | CPA>R-PIA>NECA>IB-MECA>SPA>DPMA>S-PIA>CGS21680 | (9) |
| A1 | Agonists | Rat (adipocyte, left & right atrium) &Guinea pig (ileum, left & right atrium) | | CPA≥GR79236, R-PIA≥NECA»S-PIA≥Metrifudil≥CV-1808, CGS21680 | (10) |
| A1 | Antagonist | Review of functional studies | | DPCPX>PD115199>8PT | (2) |
| A1 | Antagonist | Human | Brain | DPCPX>KFM19»Theophylline>Caffeine | (3) |
| A1 | Antagonist | Bovine | Brain | XAC>DPCPX>BWA1433=CPT>BWA844» Theophylline | (6) |
| A1 | Antagonist | Guinea pig | Brain | CPX»XAC>IBMX»DPMX>ADAC>DMPX | (7) |
| A1 | Antagonist | Rat | | CPX>XAC>8-PX=N-0861>CGS15943>8-PT>PD115199>8-p-SPX>CP66713>HTQZ>Tracaxolate>8-p-SPT>Alloxazine>Theophylline>CSC>Caffeine>DMPX | (9) |
| A1 | Antagonist | Rat | Testis | DPCPX>CPT>IBMX | (8) |

**Table S8.** A2a- and A2b- Adenosine Receptor Agonist and Antagonist profiles in the literature.

| Subtype | Ago/Ant | Species | Tissue | Profile | Reference |
|---|---|---|---|---|---|
| A2 | Agonist | Human | lung | NECA>L-PIA>D-PIA>CHA | (11) |
| A2 | Agonist | Rat | Kidney | NECA>2ClAdo>R-PIA>CHA | (12) |
| A2 | Antagonist | Human | lung | 8PT>IBMX>Theophylline | (11) |
| A2 | Antagonist | Rat | Kidney | PD115199>PD116948 | (12) |
| A2a | Agonist | Review of functional studies | | CGS21680=NECA>CV1808≥2ClAdo.R-PIA=CHA=CPA>S-PIA | (2) |
| A2a | Agonist | Human | Brain | NECA>2ClAdo>R-PIA>CPA≥S-PIA | (4) |
| A2a | Agonist | Human | Brain | CGS21680>NECA>2ClAdo>CPA | (13) |
| A2a | Agonist | Dog (coronary) & Human (platelets and neutrophils) | | CV-1808, CGS21680≥NECA>R-PIA≥Metrifudil≥CPA>GR79236, S-PIA | (10) |
| A2a | Agonist | Rat | | DPMA>NECA>CGS21680>IB-MECA>R-PIA>S-PIA>CPA>SPA | (9) |
| A2a | Agonist | Rat | Brain | CGS21680≈NECA≥CV1808 | (14) |
| A2a | Antagonist | Review of functional studies | | PD115199>DPCPX=8PT | (2) |
| A2a | Antagonist | Human | Brain | 8PT>XAC | (13) |
| A2a | Antagonist | Rat | | CGS15943>CP66713>PD115199>CSC>XAC>HTQZ>8-PX>CPX>8PT>8-p-SPX=CPT>TracazolateN-0861>8-p-SPX>CMPX>Alloxazine>Theophylline>Caffeine | (9) |
| A2a | Antagonist | Rat | Brain | CGS15943>CGS22988 | (14) |

| Subtype | Ago/Ant | Species | Tissue | Profile | Reference |
|---|---|---|---|---|---|
| A2b | Agonist | Review of functional studies | | NECA>2ClAdo>R-PIA=CHA>S-PIA≥CV1808≥CGS21680 | (2) |
| A2b | Agonist | Human | Brain | Low affinity, increase cAMP with NECA & no affect with CCPA and CGS21680 | (15) |
| A2b | Agonist | Guinea pig | Aorta | NECA>Metrifudil>R-PIA, CPA>CV-1808, GR79236≥S-PIA, CGS21680 | (10) |
| A2b | Agonist | Rat | | NECA>R-PI-NECA>2ClAdo>R-PIA>CHA>S-PIA>CGS21680 | (9) |
| A2b | Agonist | Rat | Brain | Binding ++NECA, -CGS21680, - CCPA | (16) |
| A2b | Antagonist | Review of functional studies | | DPCPX=8PT≥PD115199 | (2) |
| A2b | Antagonist | Human | Brain | Inhibited by theophylline | (15) |
| A2b | Antagonist | Rat | | CGS15943>PD115199>CP66713>XAC>CPX>8PT>Alloxazine>8-p-SPT>HTQZ>Tracazolate>CSC>DPMX>Caffeine>Theophylline | (9) |

**Table S9**. Amino acid identity, shark $A_0$ and human specialized receptors (all vs. all).

| | Shark $A_0$ | Human $A_1$ | Human $A_{2a}$ | Human $A_{2b}$ | Human $A_3$ |
|---|---|---|---|---|---|
| Shark $A_0$ | **100%** | 39% | 42% | 41% | 36% |
| Human $A_1$ | **39%** | 100% | 51% | 45% | 47% |
| Human $A_{2a}$ | **42%** | 51% | 100% | 58% | 41% |
| Human $A_{2b}$ | **41%** | 45% | 58% | 100% | 38% |
| Human $A_3$ | **36%** | 47% | 41% | 38% | 100% |
